# Facultative variation across a shallow to deep torpor spectrum in hummingbirds

**DOI:** 10.1101/2021.07.18.452827

**Authors:** A Shankar, INH Cisneros, S Thompson, CH Graham, DR Powers

## Abstract

Many small endotherms use torpor, saving energy by a controlled reduction of their body temperature and metabolic rate. Some species (e.g. arctic ground squirrels, hummingbirds) enter deep torpor, dropping their body temperatures by 23-37 °C, while others can only enter shallow torpor (e.g., pigeons, 3-10 °C reductions). However, deep torpor in mammals can increase predation risk (unless animals are in burrows or caves), inhibit immune function, and result in sleep deprivation, so even for species that can enter deep torpor, facultative shallow torpor might help balance energy savings with these potential costs. Deep torpor occurs in three avian orders. Although the literature hints that some bird species can use both shallow and deep torpor, little empirical evidence of such an avian torpor spectrum exists. We infrared imaged three hummingbird species that are known to use deep torpor, under natural temperature and light cycles, to test if they were also capable of shallow torpor. All three species used both deep and shallow torpor, often on the same night. Depending on the species, they used shallow torpor for 5-35% of the night. The presence of a bird torpor spectrum indicates a capacity for fine-scale physiological and genetic regulation of avian torpid metabolism.

## Introduction

Torpor is an energy saving strategy documented in over 200 species of birds and mammals [1]. Torpid animals save energy by lowering their metabolic rate and body temperature as their environment gets colder. Much of the relatively recent work on the metabolic torpor spectrum has focused on mammals [1–3]. A flexible physiological continuum from shallow to deep torpor seems to exist in mammals, given energetic, neurological (EEG), transcriptomic, and ecological data, as found in several ground squirrel species, marmots, and kangaroo rats [4–11]. Some bird species are known to use shallow torpor at night, while others regularly use deep torpor [1]. Though avian shallow torpor and deep torpor have separately received research attention [12–16], the differences and potential trade-offs between these states in birds are poorly studied relative to mammals. There are some hints in the literature that such a torpor spectrum exists in birds under specific conditions [in mousebirds, 17]. Though birds constitute 65% of extant endotherms, the data on avian heterothermy are sparse compared to mammalian data [18,19], as are data on this avian torpor spectrum. Exploring the range, variability, and flexibility of avian torpor can help elucidate behavioural and physiological mechanisms underlying thermoregulation, energy regulation and torpor use across vertebrates, and move us closer to understanding the evolution of homeothermy vs. heterothermy.

Heterothermic animals are often described as having species-specific minimum torpid body temperatures [between −2 and 29.6 °C, 1,20–23]. Depending on their minimum torpid body temperatures, some birds only use a ‘shallow’ form of torpor (e.g. pigeons, body temperature 28 – 36 °C; Figure 1c), while others use ‘deep’ torpor, in which body temperature is low (e.g. hummingbirds, 3 – 18 °C; Figure 1b). Of the 42 bird species reported to use daily torpor, only hummingbirds (*Trochilidae*), nightjars (*Caprimulgidae*), and one mousebird (*Coliidae*) species have minimum body temperatures colder than 20 °C; the rest use a relatively shallow form of torpor [1].

**Figure 1:**
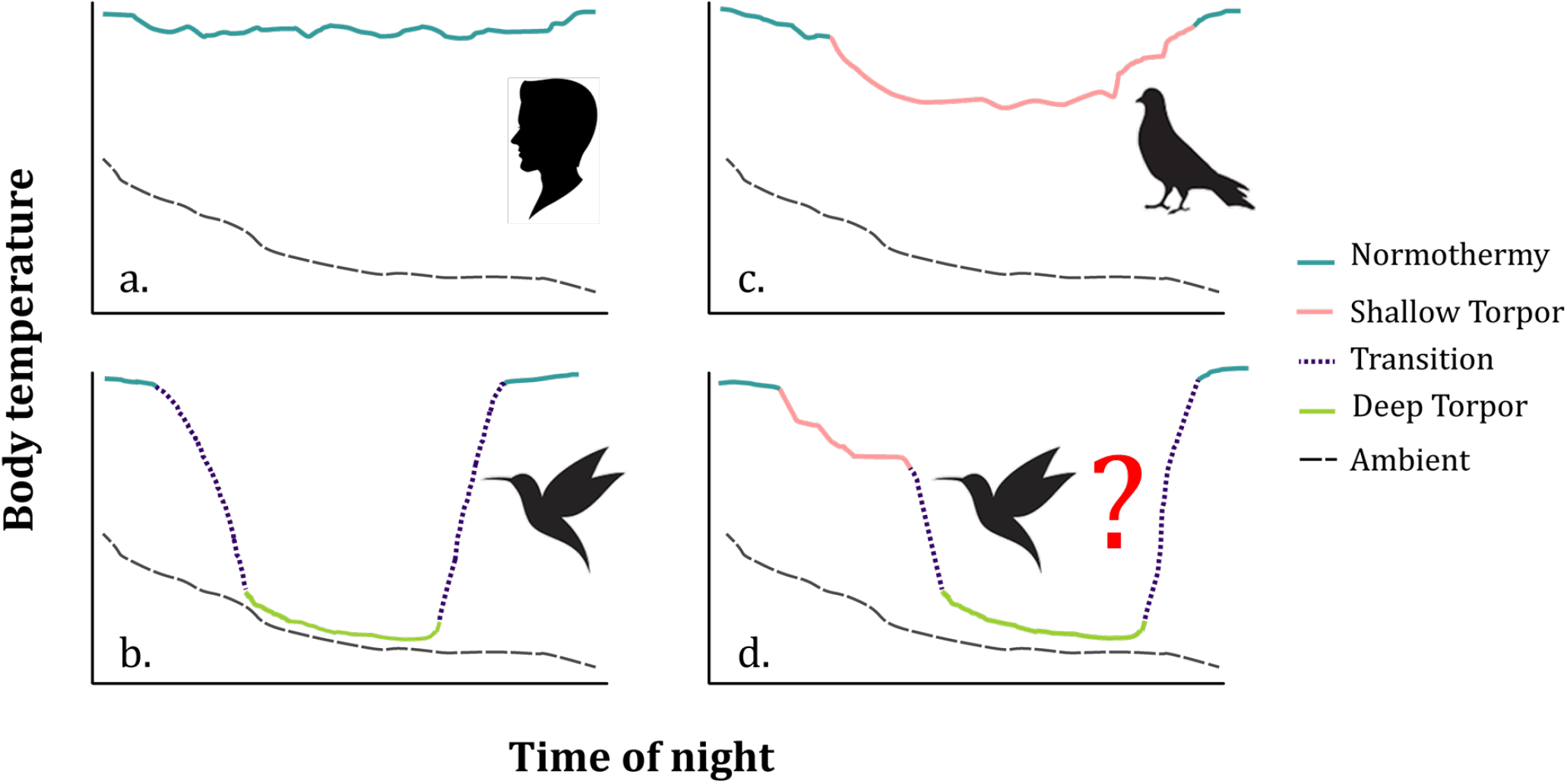
A schematic depiction of body temperature (coloured lines) relative to ambient temperature (black dashed line) at night, in sleep, shallow torpor, and deep torpor. a. A normothermic individual, with minimal circadian reductions in nighttime body temperature (e.g. humans). b. An individual starts the night normothermic, then transitions into deep torpor, where body temperature drops with ambient temperature, minimizing the difference between minimum body temperature and ambient temperature (e.g. hummingbirds). c. An individual starts the night normothermic, then transitions into ‘shallow’ torpor, potentially because the species has a very high minimum body temperature of only 4-5 °C below normothermic levels (e.g. some pigeon species). d. An individual uses a combination of normothermy, shallow, and deep torpor, at times regulating its body temperature above its minimum torpid body temperature – here we investigate the presence of such a torpor spectrum in hummingbirds.

Deep torpor likely reflects a trade-off between its benefits—an average of 60% energy savings relative to basal metabolic rates [24,25]—and potential costs such as susceptibility to predation, inability to rewarm, immune suppression, and sleep deprivation [3,26–30]. Given these trade-offs, it might be beneficial for birds that use the deepest possible form of torpor to sometimes use a shallower form, to allow moderate energy savings while minimizing some of the potential costs of deep torpor [31]. Yet in contrast to mammals, birds that are known to use deep torpor do not seem to use a shallower version of torpor by regulating their body temperatures above this minimum [1,5,15,20,32–34]. One mousebird species has been described to use both shallow and deep torpor when starved over several days, with their depth of torpor deepening as their energy stores were depleted [35,36]. However, mousebirds are thought to have diverged early in the avian phylogeny and their unusual combination (among birds) of low-quality plant diet with a relatively small body size makes them physiologically distinct in other ways. It is also possible that they display a form of ‘proto-torpor’ without the standard entry and rewarming patterns of other avian lineages [36–38]. The rarity of shallow torpor in birds that use deep torpor, and vice versa, would imply that shallow and deep torpor are mutually exclusive and relatively inflexible states. The possible existence of a torpor continuum has been hinted at in the literature [20], but evidence supporting or disproving its existence is scarce.

Hummingbirds have long been known to use deep torpor to save energy overnight, with minimum body temperatures varying from 3-22 °C [21,25,39,40]. One past study reported a shallower form of torpor in hummingbirds, but its experimental conditions may have prevented deep torpor: those birds were maintained at warm temperatures, were not free-living, and were frequently disturbed at night [41]. If hummingbirds can use both shallow and deep torpor, they are facultatively controlling their body temperature and metabolism over a broad range of torpid temperatures, despite much lower ambient temperatures. This flexibility in body temperatures is almost never described in birds, but such a capacity could contribute to hummingbirds’ ability to thrive under diverse and variable environmental conditions, from deserts to tropical forests and from sea level to the high Andes, despite their small body size and extreme metabolic demands. Previous work suggested that some larger hummingbird species had more variable metabolic rates than smaller hummingbirds [24,39], and our preliminary data from sites in the high Ecuadorian Andes has also suggested that some hummingbird species there might be using a range of shallow and deep torpor (Shankar et al., unpub).

Here we test whether hummingbirds are capable of shallow torpor by recording nighttime surface temperatures in three species sympatric at sites in Arizona (USA) where nighttime temperatures are cold enough to allow deep torpor. We know from previous work that all three species use deep torpor [16,24]. We hypothesized that these hummingbirds might facultatively use shallow torpor to balance the energy savings and physiological costs of deep torpor. Hummingbirds appear to delay torpor until they have reached some minimum threshold of energy stores [16,25,42]. We therefore expected birds to use shallow torpor in one of two ways: either exclusively with normothermy (Figure 1c), or before entering deep torpor, as a strategy to delay the onset and potential costs of deep torpor (Figure 1d). Given that hummingbirds seem to reach a minimum energetic threshold before entering deep torpor [16], we expected that once a bird entered deep torpor, it would stay in deep torpor for the remainder of that night, and then rewarm to normothermy before flying off, rather than using shallow torpor after deep torpor. We use thermal imaging to study hummingbird torpor under near-natural conditions. This study design allows us to assess torpor use under natural light and temperature cues, as well as near-natural energy stores. If hummingbirds can use shallow as well as deep torpor, it would imply that they are able to regulate their metabolism and body temperature dynamically and with great flexibility. Such physiological control in torpor would in turn imply that a broad and perhaps continuous avian metabolic torpor spectrum exists, much like in mammals.

## Methods

### Study sites and species

We studied males of three hummingbird species at the Southwestern Research Station (SWRS) in the Chiracahua mountains of Arizona (Lat: 31.9, Long: −109.2): the blue-throated mountain-gem (*Lampornis clemenciae;* 8.4g, n = 14), Rivoli’s hummingbird (*Eugenes fulgens*; 7.6g, n =12) and the black-chinned hummingbird (*Archilocus alexandri*; 2.9g, n = 7). Two blue-throated mountain-gem individuals had some bill corrugation and were likely late-stage juveniles. Within this hummingbird community, both the black-chinned and Rivoli’s hummingbirds are subordinate to blue-throated mountain-gems (i.e., with less exclusive access to floral resources) [16,43]. We collected data between June 10 – 19, 2017 and May 20 – June 7, 2018.

### Thermal imaging—nighttime surface temperatures

We captured hummingbirds using modified Hall traps at hummingbird feeders [44] within 1.5 hours before sunset, to allow them to store energy naturally through the day, but also acclimate to our experimental setup. Most birds were already banded (this is a long-term bird monitoring site), but un-banded birds were marked with a small dot of non-toxic paint on the forehead. We recorded capture mass, allowed the birds to feed *ad libitum*, and weighed them again for mass after feeding. They were then placed outdoors (individually) in five-sided acrylic chambers (either 18×17×22 cm or 46×23×46 cm), exposed to natural light and temperatures. The front face of the chamber was covered by a clear plastic sheet to prevent the bird escaping. This sheet caused the thermal reading of the bird’s surface temperatures to be up to 2 °C cooler than direct bird readings, so once the bird was observed to settle down, the plastic sheet was removed. We placed a wire grill at the base of the chamber to encourage birds to perch with their sagittal plane facing the camera, usually ensuring that recordings included a direct view of the bird’s eye. Recordings without this view were excluded from analyses.

Bird eye surface temperatures seem to closely reflect internal physiological state (e.g. body condition), from recent work in blue tits [45]. Hummingbirds have low feather density around the eye, so skin eye temperature patterns should closely reflect the patterns of core body temperature, minimizing the confounding effects of feather insulation, unlike in larger animals for which skin and core temperatures might vary because of reduced peripheral blood flow in torpor [15,46–49]. The traditional rules of endotherm thermoneutrality and body temperature do not seem to apply in hummingbirds because of their small size and status as ‘micro-endotherms’; they do not seem to maintain steady-state thermal equilibrium as a result [50]. . Powers et al. [51] used thermal imaging to measure heat dissipation areas in hovering hummingbirds during the day in three species. They found that across all three species, eye surface temperature remained relatively constant across a range of ambient temperatures, with an intercept of 32–33 °C [Figure 2 in 51]. Although this is lower than core body temperature, it is consistent with what is observed in individuals from these same species that are clearly normothermic. This supports our measurements of 32 °C being a common resting normothermic body temperature. A recent study of hummingbird body temperature in torpor across six species showed that many individuals maintained normothermy at body (cloacal) temperatures below 40 °C [39]. Additionally, there seems to be high concordance between eye surface temperatures and metabolic rates as measured in ruby-throated hummingbirds (Erich Eberts, unpub.).

**Figure 2:**
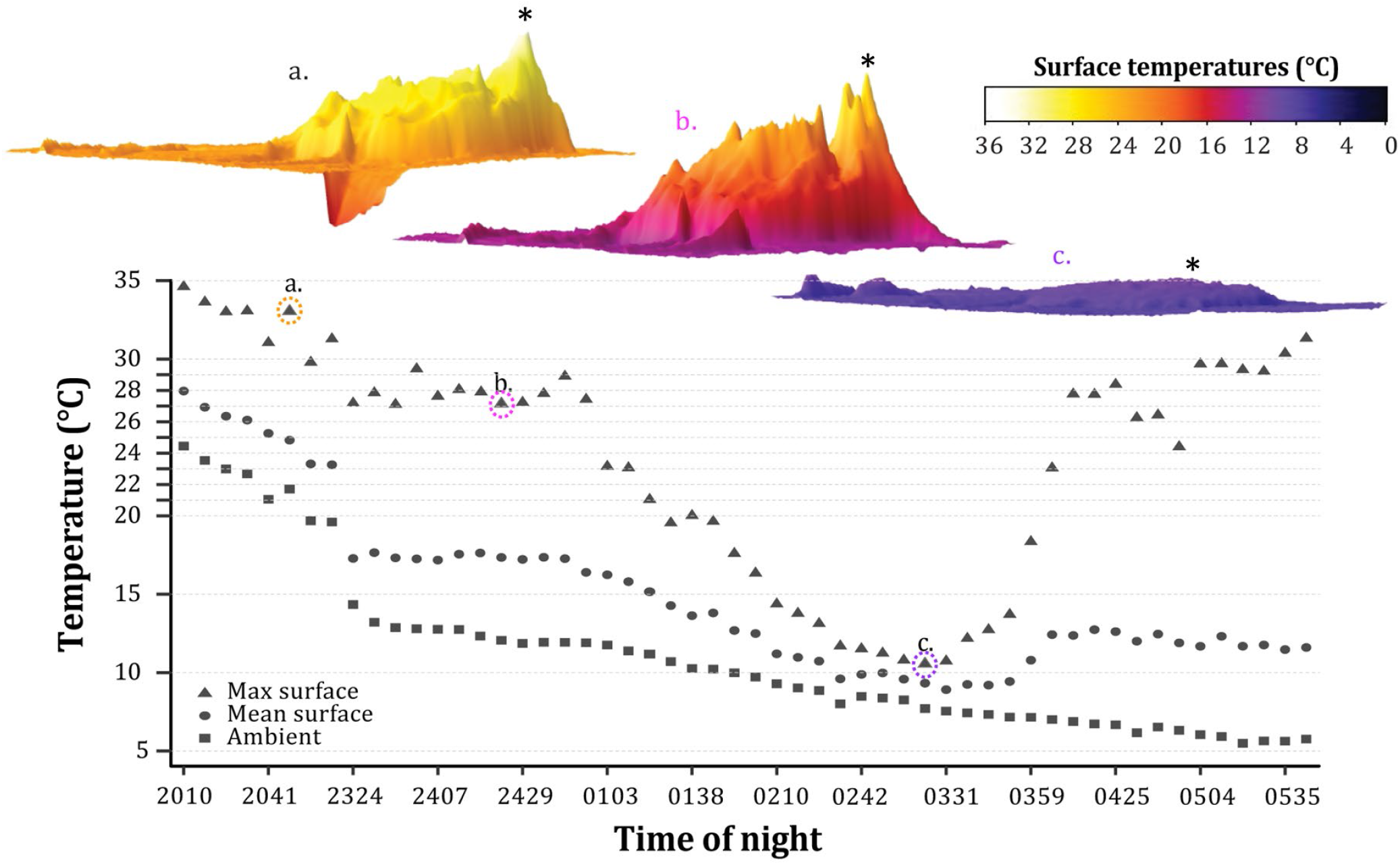
A Rivoli’s hummingbird using all four metabolic states (normothermy, shallow torpor, transition and deep torpor). Top: 3D plots of the surface temperature of the bird in normothermy, shallow torpor, and deep torpor, aligned with the tail-beak axis along the x-axis. Asterisks indicate the location of the eye. See Supplementary Video SV1 for perspective on the 3D plots. a. Normothermic: surface temperatures peak at 35 °C near the eye; mean surface temperature is 25 °C. b. Shallow: surface temperatures peak around the eye at 27 °C, followed by a drop in temperature and steady, much lower, surface temperatures over the rest of the body (17 °C), and then a steady drop towards the tail. c. Torpor: The entire surface of the bird is cold, peaking around the eye at 11 °C.

We used a FLIR SC6701 infrared video camera (640 × 480 pixel resolution, sampling at 300 Hz; accurate to 1 °C at measured temperatures) to record surface temperatures of hummingbirds. We assumed emissivity was 0.95 across all surfaces of the hummingbird [47,52]. We monitored birds continuously through the night, and sampled surface temperatures by recording 10 seconds of 30Hz video approximately every 10 minutes, using ResearchIR (FLIR, Inc.). From one frame per recording, a region including the bird and a slight buffer to include ambient temperatures was marked as a region of interest and exported to csv files for analysis in R [v.3.5.1; 53]. From each exported region of interest, we extracted maximum surface temperature (in Celsius) and mean surface temperature of the bird, as well as minimum temperature (our proxy for ambient temperature). We verified that maximum surface temperatures corresponded with maximum eye surface temperatures, and validated outliers in temperature measurements to ensure that they were reliable measurements. We also exported entire single-frame images from selected recordings and used ImageJ (NIH) to construct 3D images to assess how surface temperatures changed over the entire surface of the bird.

### Ambient temperature

We used minimum temperatures from thermal image regions of interest as an estimate of ambient temperatures, verifying that these closely matched independent ambient temperature measurements from iButtons (Maxim Integrated DS1921) or thermocouples (Cu-Cn type-T, recorded on a TC-1000; Sable Systems). The FLIR camera was factory calibrated and verified by imaging a surface of a known temperature. Thermocouples and iButtons were calibrated by using a Percival (model I-35LV, Percival Scientific, Inc.) at controlled temperature steps, and checked against a thermometer traceable to the National Institute of Standards and Technology.

### Thermal categories

We assigned bird surface temperature measurements at each time point to one of four categories: normothermy, shallow torpor, transition to and from deep torpor, or deep torpor. We defined these categories using individually assigned thresholds for each bird. We used eye surface temperatures of the bird once it had settled, but its eyes were still open, to define resting normothermic temperatures. Once the eyes were closed, we considered the bird asleep [54]. If eye surface temperatures dropped more than 2 °C below these resting temperatures, we classified the birds into one of the other three categories based on 1. rate of temperature change (stable, slow change, rapid change), and 2. magnitude of decrease of eye surface temperature below normothermic temperature, and above ambient temperature.

Birds were considered in shallow torpor if they dropped more than 2 °C but less than 20 °C below their resting temperature (but were still above ambient temperature), and maintained that temperature for more than 10 minutes (stable temperatures). Measurements were assigned to the transition category if they dropped or increased rapidly between normothermy and deep torpor, or between shallow and deep torpor (i.e. transitions were defined by rapid, large temperature changes; average rate of change ± s.e. of 0.45 ± 0.06 °C/min, up to 3 °C/min; see Figure S2 for details). Birds were considered to be in deep torpor if eye surface temperature was close to ambient temperature, or if it was maintained below 20 °C without dropping any lower (stable, low temperatures), for an extended period [highest reported hummingbird torpid body temperature is 22 °C, 21,55].

### Surface temperatures models

Normothermy, shallow torpor and deep torpor could be distinguished by the relationship between surface (response) and ambient (predictor) temperature. While a normothermic homeotherm can maintain a relatively stable body or surface temperature over a large range of ambient temperatures, in deep torpor the body and surface temperatures become a positive function of ambient temperature. Therefore, we would expect deep torpor to have a steep slope and a very low intercept. If shallow torpor exists, then the slopes for normothermy and shallow torpor should be similar and low, while their intercepts should vary (shallow torpor lower than normothermy). Additionally, we expected species to use these torpor categories differently, and expected mass to negatively influence torpor use [birds with greater energy stores should use torpor less, 16].

To estimate regression equations of surface temperature as a function of ambient temperature for each of the four thermal categories (normothermy, shallow torpor, transition, deep torpor) we used linear mixed effects models [56] using the ‘nlme’ package in R [57]. A mixed effects model is appropriate because the response (surface) can be modelled as a function of various data types; in this case both continuous fixed effects (ambient temperature, thermal categories, mass, species, and year), as well as random effects (categories nested within individuals) and an autocorrelation term were incorporated. To first test the effect of ambient temperature on surface temperature, we ran a simple linear model of surface temperature (*T*_*S*_) as a function of ambient temperature (*T*_*a*_). This model only explained 15% of the variation in surface temperatures, and we therefore ran an ‘lme’ linear mixed effects model of surface temperature as a function of ambient temperature. We included mass as a continuous fixed covariate; thermal category (normothermy, shallow torpor, etc.), species, and year as discrete fixed covariates, and categories within individuals as a random covariate. We included interaction terms between category and both ambient temperature and species. We also included an autocorrelation term (‘CorAR1’; see Supplement 1 for model details):

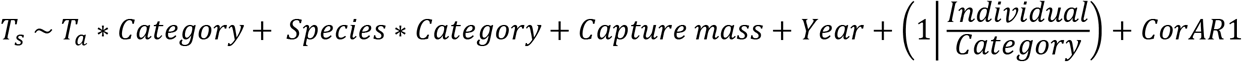

### Frequency of thermal category use

To estimate the proportion of time that each species spent in each of the four categories, we calculated the proportion of the night spent in each thermal category for every individual. We then modelled the percentage of the night spent in each category per species. We ran thermal category and species as interacting terms because we expected them to have interactive effects. The model was:

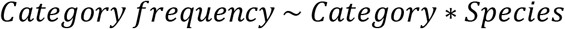

We ran a generalized linear model, fitting a negative binomial distribution to the data [58]. We first ran Poisson and quasipoisson models, but both were overdispersed (see Supplement 1 and Table S1). We therefore fit a negative binomial which was much better fit than the others (Table S2). We used the ‘glm.nb’ function in the MASS package in R to run this model [53,58,59].

## Results

### Ambient temperatures

Ambient temperatures usually declined steadily over the course of the night (1930h and 550h, e.g. in Figure 2). In 2017, temperatures were an average ± s.d. of 13 ± 4.6 °C (range 3 to 23 °C), and in 2018 they were 11 ± 5.7 °C (range −1 to 24 °C). Most nights ranged between 5 – 20 °C (mean 12 °C), except for one particularly cold night with ambient temperatures between −1 and 14 °C (May 20, 2018), and one especially warm night (June 5, 2018) during which ambient temperatures ranged between 15 – 25 °C.

### Nighttime surface temperatures

The surface temperatures of normothermic birds and birds in shallow torpor peaked near the eye and decreased from the eye towards the tail (Figures 2 and 3). Birds in deep torpor were evenly cold. Nighttime eye surface temperature varied overall between 5.9-38 °C (Supplement 2). Active birds at the beginning of the night had normothermic temperatures ranging between 31-38 °C. This wide range included birds that were hovering and birds at rest. When they settled down, normothermic temperatures usually stabilized (when the bird was resting with eyes open) at 31 °C, so we usually considered minimum normothermic resting surface temperatures to be around 31 °C. In some cases, birds stabilized at 29 °C, at both the start and end of the night, with minimal fluctuation; in these cases we set the resting normothermic threshold to be 29 °C. Maximum eye surface temperatures ranged from 29-38 °C in normothermy, to 19.5–29 °C in shallow torpor, and 5.9–24.1 °C in deep torpor. Categories varied slightly across individuals because we assigned category thresholds per bird based on its surface temperature patterns relative to resting and ambient temperatures, and based on the rate of temperature change.

**Figure 3:**
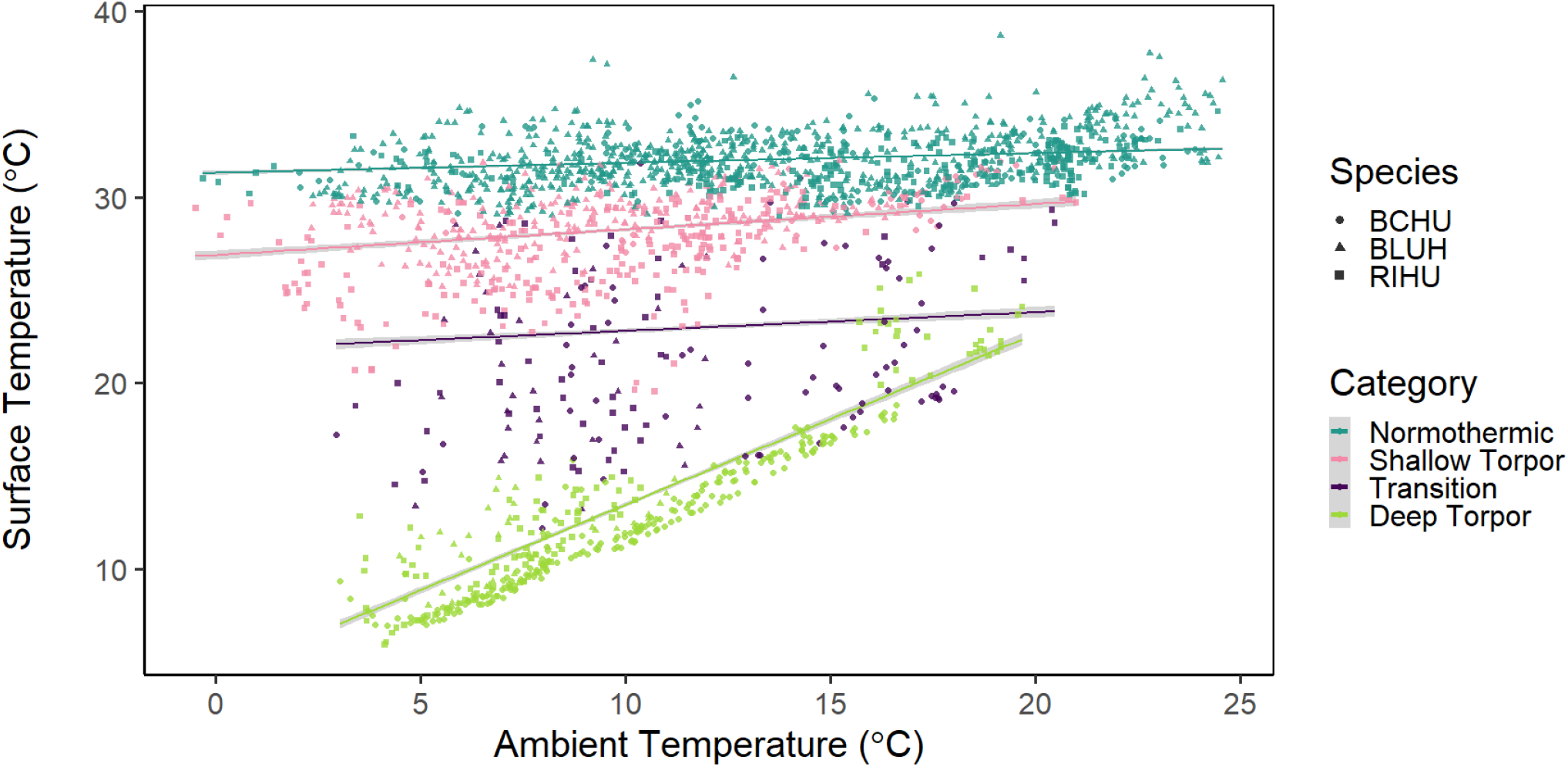
Predicted model fit from the linear mixed effects model of maximum surface temperatures (eye temperatures) as a function of ambient temperatures, coloured by category. As we predicted, deep torpor had a steep slope and low intercept, while shallow torpor and normothermy had similar low slopes and high intercepts (see Tables S3 and S4 for regression coefficients).

### Surface temperature model results

The full model for surface temperature (where the slopes and intercepts vary by category and species) allowed us to identify and quantify the various thermal categories, including shallow torpor (Figure 3, Table S3 and Table S4). Mass did not seem to have a large effect surface temperature given the other factors, but year did.

The normothermic and shallow torpor categories had similar, very low, slopes (0.11), while the normothermy intercept was about 4 °C higher than the shallow torpor intercept. A 4 °C drop from normothermy has previously been categorized as being a form of torpor [1]. In contrast with these thermoregulating states, hummingbirds largely thermoconform in deep torpor (down to the ambient temperatures we measured them at, which were all above their minimum body temperatures in deep torpor). In deep torpor, their surface temperatures closely tied to ambient temperature (slope of 0.85) and a low intercept about 20 °C lower than the normothermy intercept). The transition category is a non-equilibrial physiological state, with an intermediate intercept 17 °C lower than normothermy.

### Frequency of thermal category use

Shallow torpor was used by all species, but at varying frequencies (Figure 4, Figure S1). Of the 33 individuals we studied, all 33 were normothermic for part of the night; 24 used shallow torpor for part of the night; 17 transitioned between deep torpor and normothermy, and 17 used deep torpor (Figure 4a).

**Figure 4:**
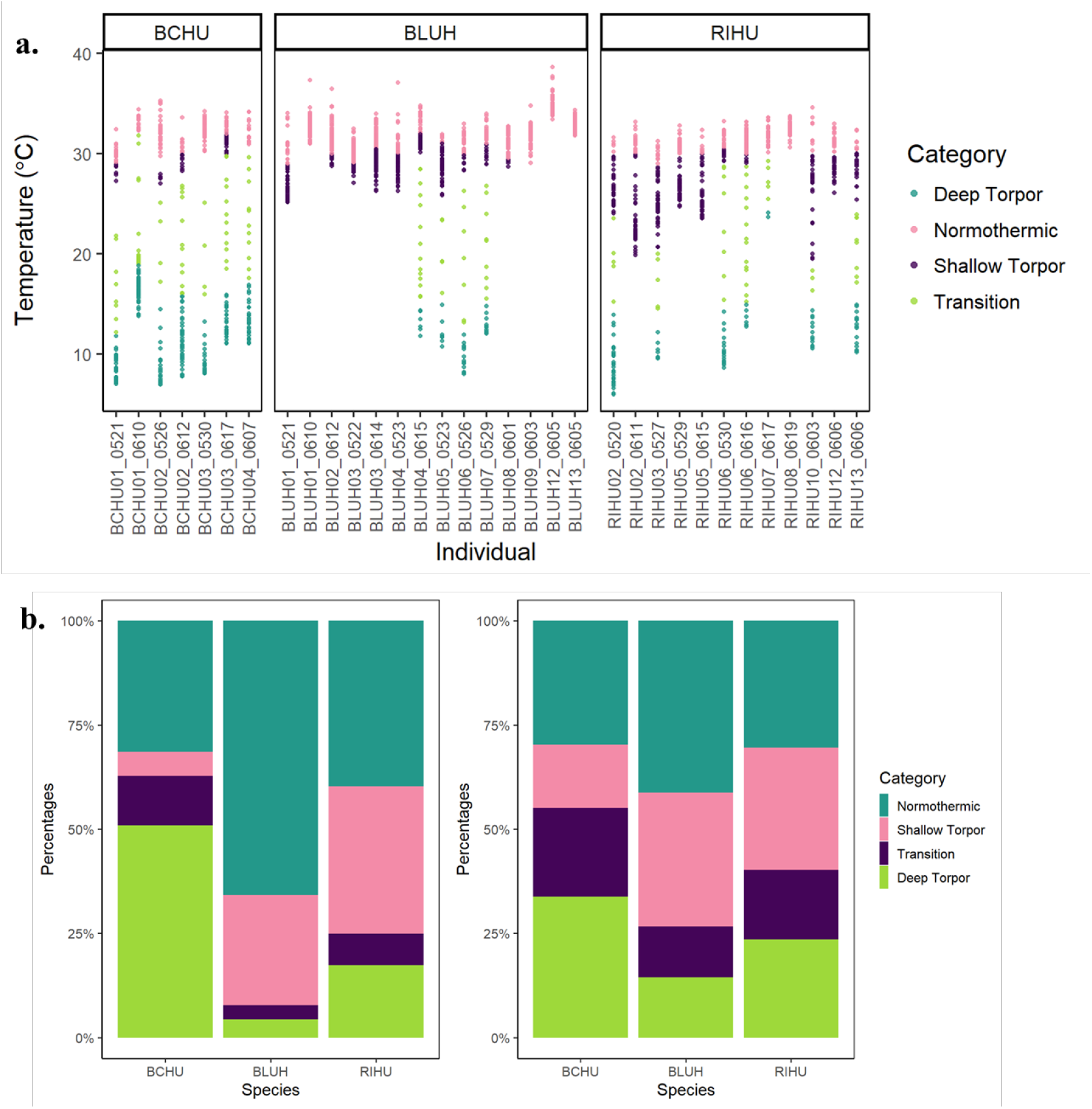
**a**. Distribution of values recorded per category (colours), per individual. BCHU = black-chinned hummingbird; BLUH = blue-throated mountain-gem; RIHU = Rivoli’s hummingbird. **b.** The relative percentages of time a species spent over all nights studied (1900h-0559h) in each of the four categories: normothermic, shallow torpor, transition, torpor. BCHU = black-chinned hummingbird; BLUH = blue-throated mountain-gem; RIHU = Rivoli’s hummingbird. Left: Percentages calculated using raw data. Right: Model estimates from the glm model (Category frequency ~ Category*Species - 1), of the relative time per category per species, presented as percentages.

All seven black-chinned hummingbirds (BCHU) used deep torpor, for an average of 49% of the night, while only three of these individuals used shallow torpor, for an average of 5% per night. BCHU spent 34% of the night on average in normothermy, and 12% in transition. The 14 blue-throated mountain-gems (BLUH) remained largely normothermic (67% of the total nighttime), and used shallow torpor an average of 25% of the night, with four individuals remaining normothermic all night, 10 individuals using shallow torpor for at least some time and only four using deep torpor (4% of the night), with 3% of the night spent on transitioning on average. The 12 Rivoli’s hummingbirds (RIHU) were the most variable in their use of the various metabolic states, with 10 individuals using shallow torpor (33% of total nighttime), six individuals using all four categories, four using normothermy and shallow torpor, one individual remaining normothermic all night, and one individual using all categories except shallow torpor. The RIHU individuals spent an average of 43% of the night in normothermy, 8% in transition, and 17% in deep torpor.

The model of thermal category frequency across species showed that there were overall clear differences in the thermal categories across all species. There were also species-specific differences in the use of normothermy, shallow torpor, deep torpor and the transition categories (Figure 4b and Table S3). The model resulted in estimates that were more evenly distributed than the raw data (the two panels on Figure 4b), but the differences between species were still clear.

The two blue-throated mountain-gems that we studied on an especially warm night maintained high surface temperatures all night (33-35 °C). The two blue-throated mountain-gem individuals that could have been late-stage juveniles appeared to behave similarly to adults: one used all four categories, and the other used only normothermy and shallow torpor. Contrary to our expectations, several Rivoli’s hummingbirds and a few blue-throated mountain-gems used shallow torpor for 1-2 hours after coming out of deep torpor, while the black-chinned hummingbirds never used shallow torpor after deep torpor.

## Discussion

We describe and quantify the novel use of shallow torpor in birds that are known to use deep torpor. Similar to mammals, and in contrast with previous studies that either describe birds as using shallow torpor or deep torpor, hummingbirds appear capable of using both. Hummingbirds in shallow torpor appear to thermoregulate to maintain surface temperatures below normothermy. In contrast, hummingbirds in deep torpor largely thermoconform to ambient temperatures. The intermediate shallow state could serve to balance nighttime energy savings with the potential ecological and physiological costs of deep torpor. Reflecting what previous studies have found [16], birds with larger energy stores seem to have greater flexibility in avoiding deep torpor. The two larger species in our study used normothermy and shallow torpor for a greater proportion of the night than the smaller species. Our minimally invasive study design allowed us to thermally image hummingbirds under near-natural temperature cycles, without disturbing or touching the birds through the night, and allowed us to discover a new level of flexibility in hummingbirds’ management of their nighttime energetic needs.

All three species used all metabolic categories, but unequally. Rivoli’s hummingbirds used shallow torpor the most, followed by blue-throated mountain-gems. The small black-chinned hummingbirds used shallow torpor the least. It therefore appears that black-chinned hummingbirds, the smallest of the three study species, might have the least flexibility in managing their nighttime energy budget, while blue-throated mountain-gems, the large territorial species, have the most flexibility. Individuals of the two larger species appeared to have more flexibility in regulating their nighttime body temperature, commonly using shallow torpor or a combination of shallow and deep torpor. The more limited use of deep torpor in these two species is consistent with previous findings that these species tend to avoid deep torpor [16,24].

Our results support the argument that there must be either physiological or ecological costs of deep torpor [1], because hummingbirds that are clearly capable of using deep torpor sometimes use shallow torpor or avoid torpor altogether. Using shallow torpor rather than deep torpor could be especially beneficial in three scenarios. First, deep torpor in mammals (especially hibernation) is usually considered helpful in avoiding predation because torpid animals are less conspicuous to predators; but these animals are usually hidden in hibernacula or dens [1,60]. Torpid birds in trees, in contrast, might be more conspicuous, making shallow torpor more efficient than deep torpor in allowing them to respond to potential predators [30]. Hummingbirds in shallow torpor could afford quicker rewarming times (< 5 minutes), and quicker responses to predators or other external stimuli, relative to deep torpor for which rewarming to normothermy takes an average of 20-30 minutes [24,61]. Second, at least in mammals, the physiological costs of torpor include rewarming costs, immune suppression [27,28], increased oxidative stress [62], and potential sleep deprivation [3,63]. There are hints that daily heterothermic mammals (Djungarian hamsters, *Phodopus sungorus*; 26g) enter a euthermic state after torpor to recover from sleep deprivation [26,63]. Shallow torpor would allow higher levels of metabolic function than deep torpor, perhaps facilitating some of the restorative functions of sleep, immunity, and lowered oxidative stress. Though avian sleep has been studied to some extent [13], little is known about the physiological basis for torpor vs. sleep in birds. Third, for nesting birds that need to keep their nest warm, shallow torpor could help balance the birds’ need to maintain energy balance with the need to supply heat to their eggs or chicks. Nesting hummingbirds have been found to generally avoid torpor (with exceptions when energy stores seemed to be low); but the use of shallow torpor by nesting birds has not been evaluated [64–66]. If deep torpor had no ecological or physiological consequences, hummingbirds would likely maximize torpor use, or remain in deep torpor for the duration of the night after entering torpor. Instead, some individuals used shallow torpor not just before a deep torpor bout as we predicted, but after emerging from a deep torpor bout, indicating that they may be trying to save energy but also balance these energy savings with the potential costs of deep torpor.

Here we identified four metabolic categories in hummingbirds: two thermoregulatory categories—normothermy and shallow torpor; a thermoconforming category—deep torpor; and the transition between deep torpor and the other categories. In normothermy and shallow torpor the animal actively thermoregulates to maintain a constant body temperature across a range of ambient temperatures. Based on the similar surface temperature slopes of normothermy and shallow torpor (Figure 3), and the rapid transitions between normothermy and shallow torpor that we often observed, these two states seem metabolically continuous in hummingbirds [67]. Shallow torpor, as defined here, could potentially be a metabolically inhibited form of normothermic sleep, but it is unclear whether the shallow torpor and deep torpor we report are on a similar metabolic spectrum, and we were unable to definitely distinguish sleep using only body temperature measurements. Multiple lines of evidence, especially in ground squirrels and pocket mice, from electroencephalograms (EEGs), measurements of brain temperature, and metabolic rates indicate that mammals slow their metabolism continuously down from sleep into torpor, and regulate their body temperatures variably above minimum body temperature [1,5,11,31,68]. Thus, though sleep and torpor appear to be on a continuous spectrum in mammals, we are yet to confirm if they are on a continuum in birds.

Given that hummingbirds can regulate between shallow and deep torpor, the biological relevance of minimum body temperature measurements must be assessed. Recent work with high-elevation Andean hummingbirds found that minimum torpid body temperature showed a phylogenetic signal, indicating that minimum torpid body temperature, at least at very cold sites, is evolutionarily conserved [39]. Shallow torpor can occur either because a bird’s minimum possible torpid body temperature is relatively high (i.e., it does not have the capacity for deeper torpor), or when a bird regulates at a high, sub-normothermic, body temperature despite its minimum torpid body temperature being much lower (e.g. 15 °C, indicating that it is capable of deep torpor; Figure 1c). These two shallow torpor scenarios are indistinguishable (as in Figure 1c) unless the species’ “true” minimum body temperature is known. If a bird regulates at a body temperature above its minimum, even though ambient temperatures were lower, these measurements might appear to be minimum body temperature measurements although they are not. In Rivoli’s hummingbirds, for instance, we found that eye surface temperatures went as low as 5.9 °C. However, Rivoli’s individuals in the laboratory were previously reported to regulate their minimum body temperature at 12 °C despite ambient temperatures going lower [14]. Such a large disparity in birds in deep torpor is unlikely to be due to differences between core and skin temperatures, and could either indicate intra-specific differences in minimum body temperature, or that the birds in the previous study were using a shallower form of torpor. This disparity could also be caused by birds reducing their blood circulation around the eye during torpor, but such regional variation seems unlikely given the small size of hummingbirds and the even distribution of low surface temperatures we observed in torpid birds. Minimum body temperature may therefore be lower than has been reported in some species. Currently minimum body temperature across all hummingbirds is thought to vary from 3–22 °C [39,40,55]. But if some of the hummingbirds measured were using shallow rather than their deepest possible torpor, the range of hummingbirds’ true minimum body temperatures would be narrower or lower. Additionally, torpor studies in hummingbirds are often conducted in laboratory conditions, which could alter torpor responses [1,20].

We propose three reasons for why this form of shallow torpor in birds has so rarely been detected [35]. First, small drops in oxygen consumption or body temperature might have been overlooked. Second, most studies of bird torpor are either done under controlled laboratory conditions, or involve handling the birds many times at night to record body temperature. Birds in captivity are often overweight and have to be starved to enter torpor [69]. Laboratory torpor studies conducted at controlled temperature steps might have pre-empted the use of shallow torpor, because shallow torpor is presumably a fine-scale response to energetic state and environmental conditions, and controlled temperature steps or repeated handling might not elicit the same physiological responses as natural decreases in nighttime temperature would [70]. Third, birds in laboratory settings are known to show altered torpor use relative to free birds: free-living animals often use torpor more frequently, and drop to a lower body temperature in torpor than laboratory animals [reviewed in 1,20,71]. Taken together, under relatively predictable natural temperature patterns, hummingbirds might be able to use intermediate torpor states more often, while in the laboratory, low temperatures, the factor most often tested, might cause the bird to either stay awake or drop into deep torpor if energetically necessary.

At the whole-animal level, the next step in understanding avian torpor would be to combine respirometry, thermal measurements and measurements of breathing or heart rates while keeping in mind the possible existence of shallow torpor. These measures have been found to sometimes be uncoupled in torpor [72,73], and therefore studying whether they vary together in shallow torpor would be the first step in identifying the physiological differences between sleep, shallow torpor and deep torpor. A promising future avenue for research would be to investigate which metabolic and genetic pathways shut down at different temperatures in hummingbird torpor. It remains to be seen if other hummingbird and bird species that use deep torpor are also capable of shallower torpor, or if such control over their torpid metabolism is unique to these two hummingbird clades. Our data indicate that these hummingbird species in a temperate environment with cold ambient temperatures often use shallow torpor; it is therefore possible that tropical species at sites with high ambient temperatures might be doing the same.

## Supporting information

Metadata for all datasets

Summarized and formatted version of data used for most plots and models

Bird masses of all individuals

Percent time spent in each thermal category

Thresholds for each category per individual bird

Raw data processed and summarized from rds files

## Author contributions

AS, DRP, CHG and INHC were involved in study conception, design and obtaining funding. AS, INHC, ST and DRP collected data; AS analysed the data and wrote the manuscript, CHG and DRP provided major comments and revisions.

## Acknowledgements

We thank Alexis Stark for help with data collection; Liliana M. Dávalos for discussions, statistical analyses, and comments on the manuscript; Benjamin Van Doren for help with statistical analyses; Erich Eberts and Elise Lauterbur for useful discussions; and Jeffrey Levinton for helpful comments on the manuscript. We thank our funders for their generous support: NASA (grant NNX11AO28G to CHG, SJ Goetz, SM Wethington, and DRP), the Tinker Foundation, National Geographic Society (9506-14, AS), the American Philosophical Society (AS), ERC-2017-ADG number 787638 (CHG), the Swiss Federal Research Institute (WSL) for funding writing and analysis visits (AS, DRP), a George Fox University Richter Scholar grant (ST), and a George Fox University Faculty Development Grant (GFU2014G02, DRP). CHG thanks the Swiss National Science Foundation (SNF) - No 173342 and European Research Council (ERC) under the European Union’s Horizon 2020 research and innovation program - No 787638. We would also like to thank all the contributors to two crowd-funded experiment.com grants (https://tinyurl.com/rknxz8u and https://tinyurl.com/vhbt4ke). We have no conflicts of interests to declare.

## Data and Code accessibility

The R scripts used to run the analyses are available on Github: https://github.com/nushiamme/TorporShallowDeep. The data associated with this manuscript are available as supplementary data files.

