## Supplement 2 for "Facultative variation across a shallow to deep torpor spectrum in hummingbirds"

Supplement 2_Individuals Overnight

Anusha Shankar

5/25/2021

```
## Registered S3 method overwritten by 'pryr':
##   method      from
##   print.bytes Rcpp
## For best results, restart R session and update pander using devtools:: or remotes::install_github('rapporter/pander')
## [[1]]
```

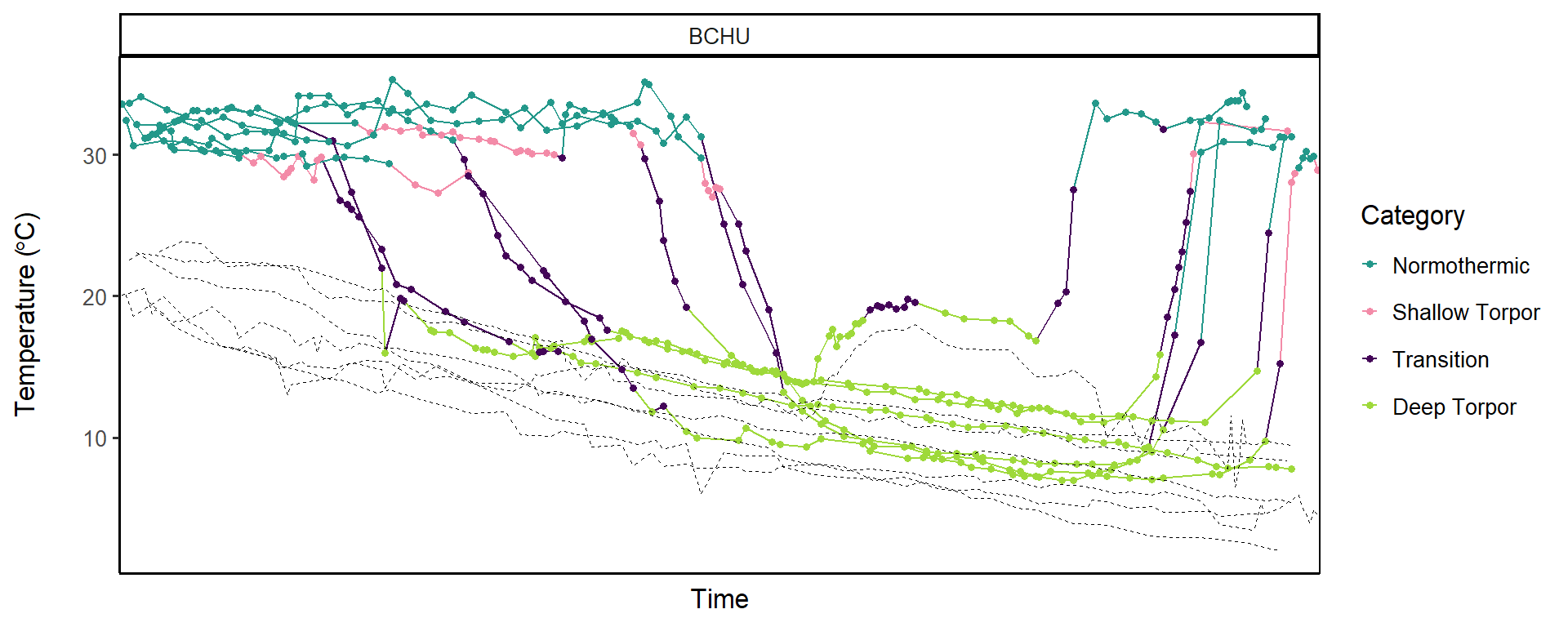

```
##
## [[2]]
```

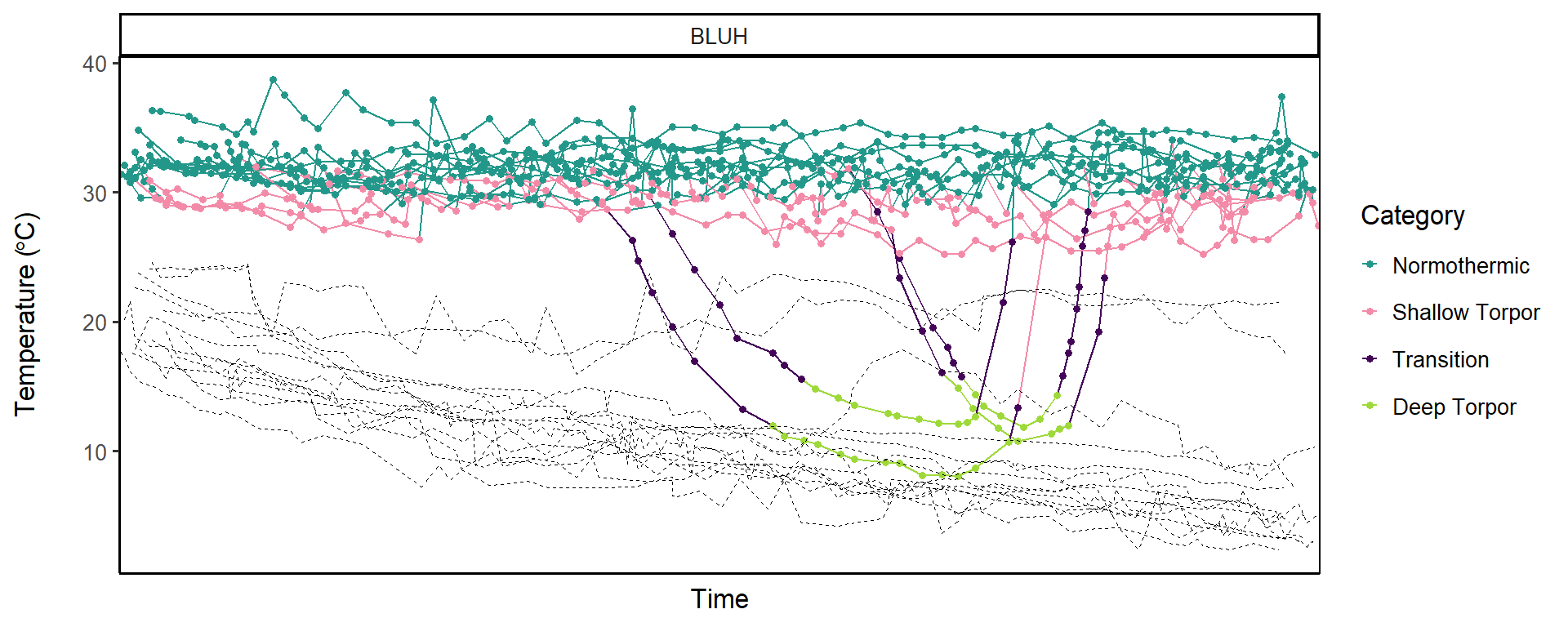

```
##
## [[3]]
```

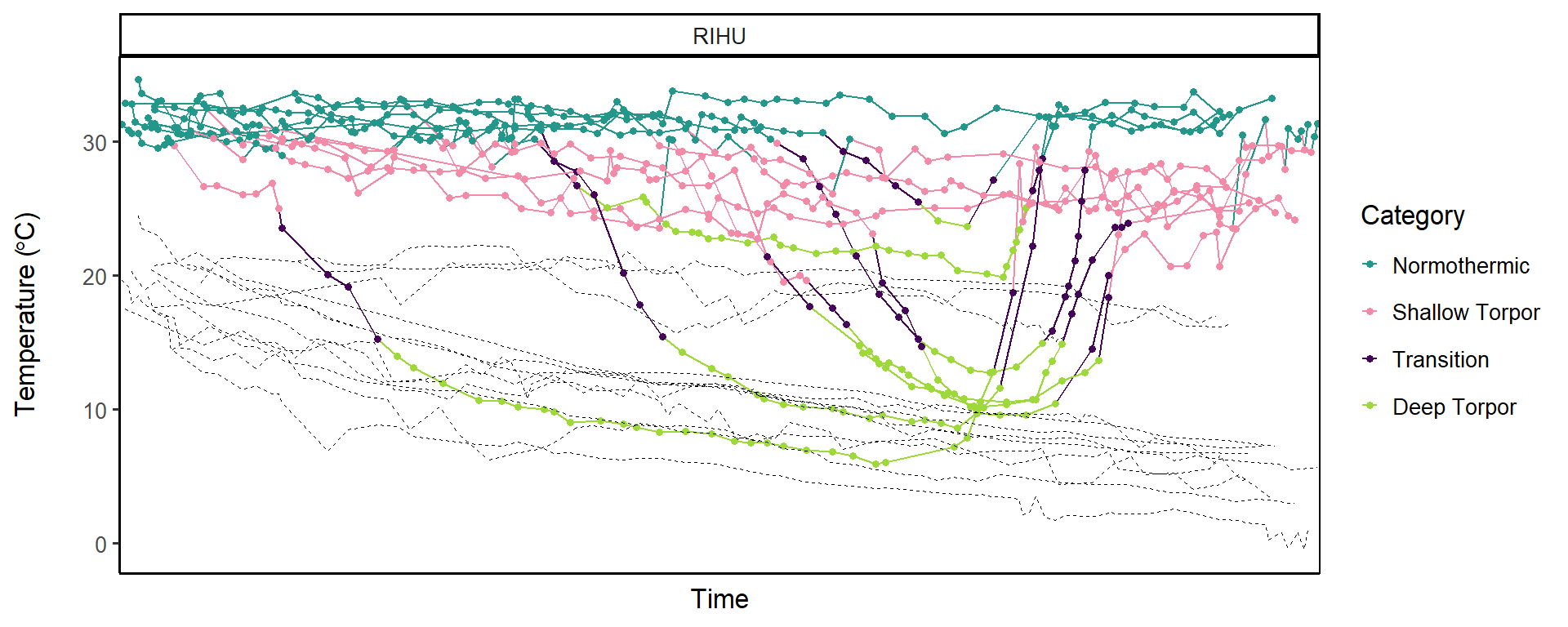

Just RIHU
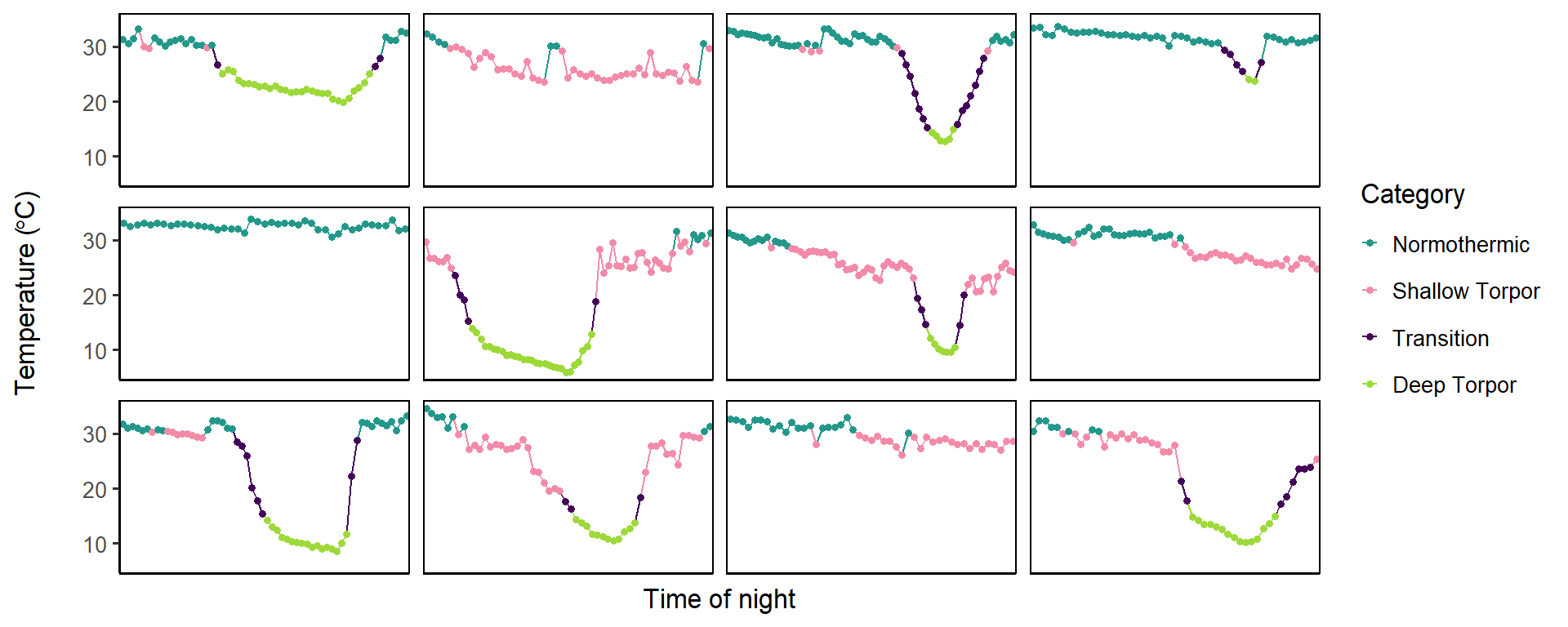

Just BLUH
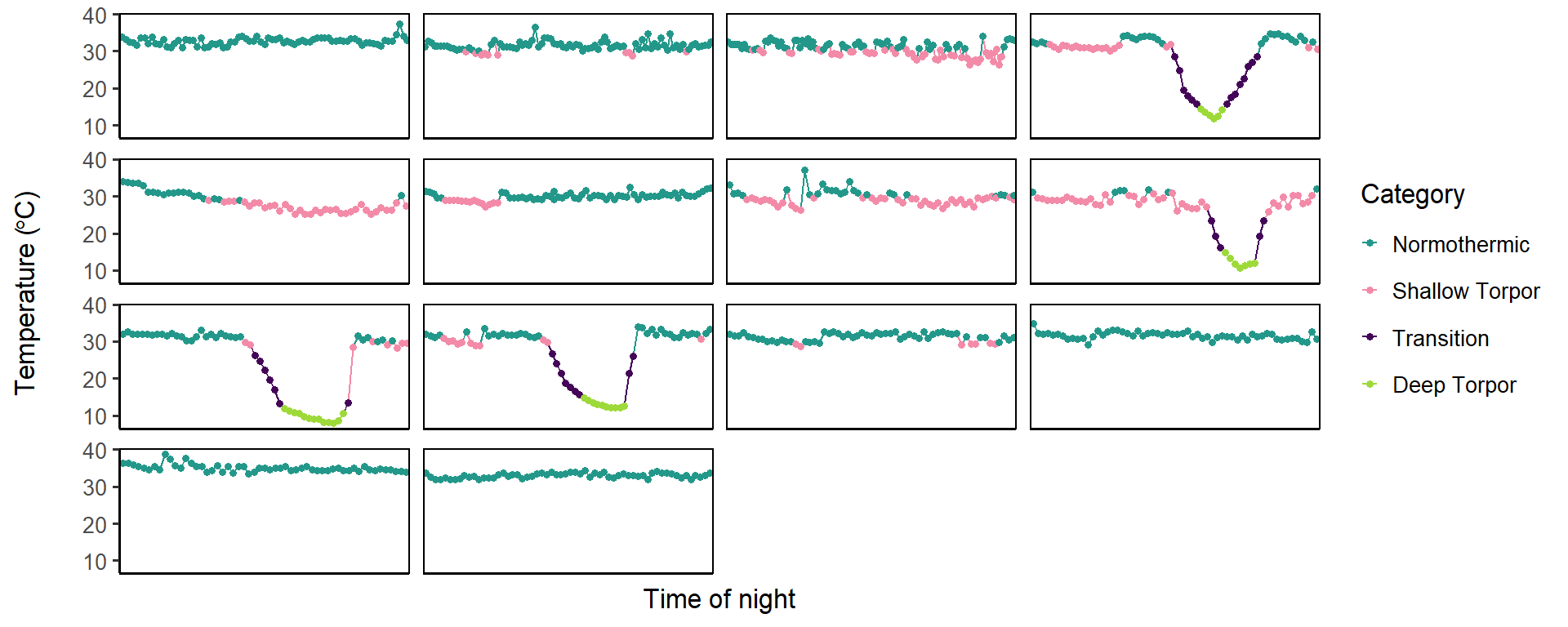

Just BCHU
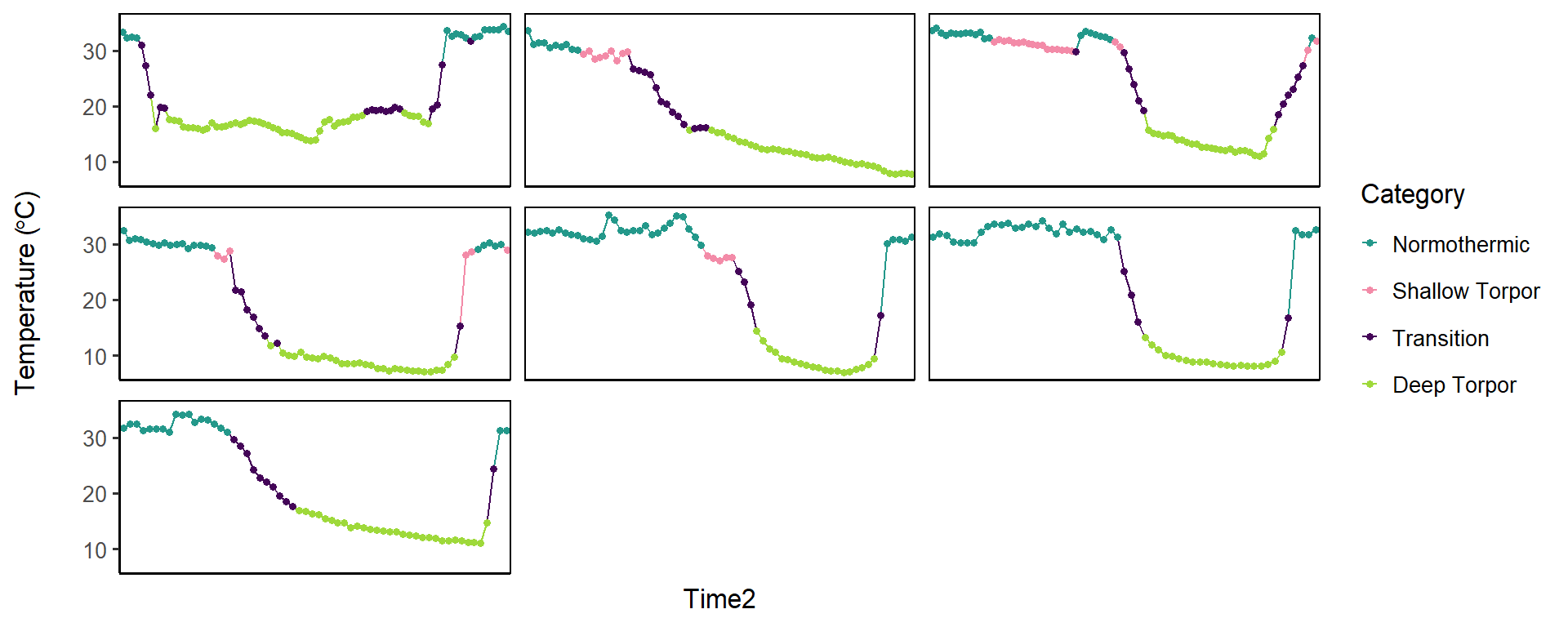

All individuals, one plot each

\## [[1]]

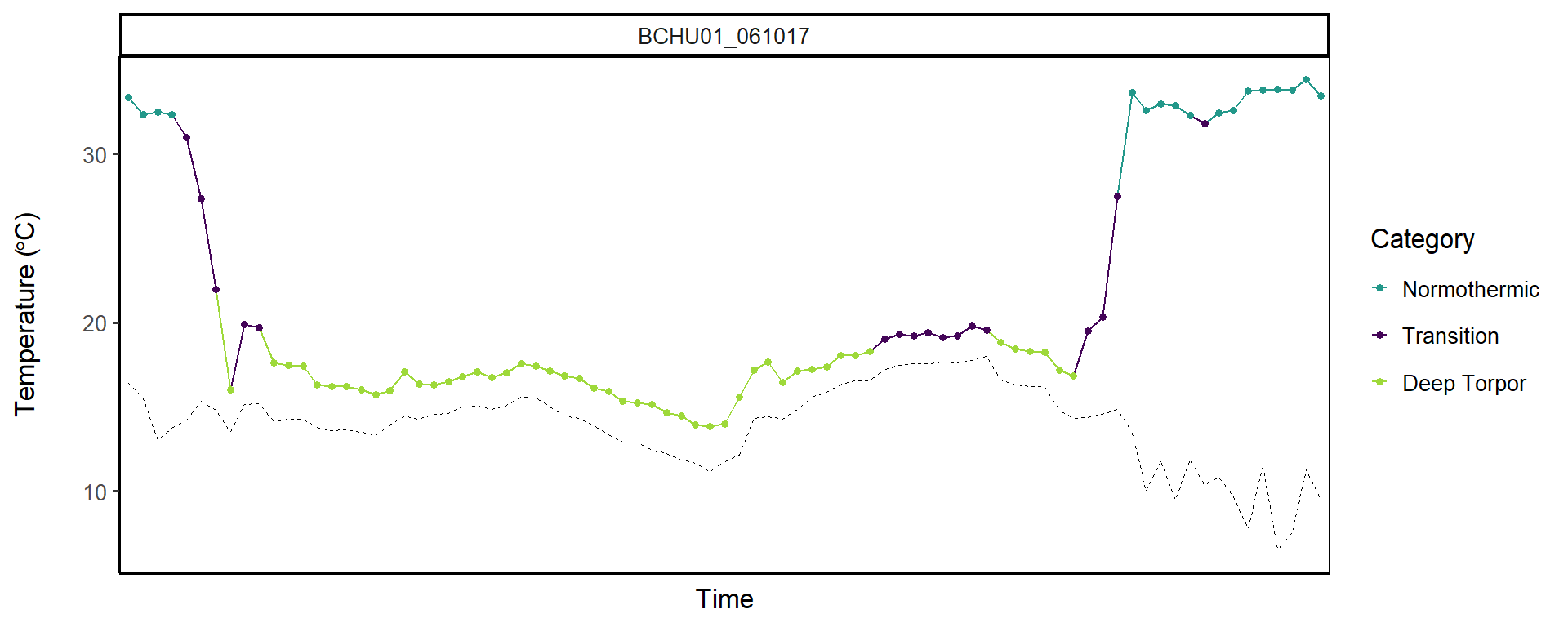

```
##
## [[2]]
```

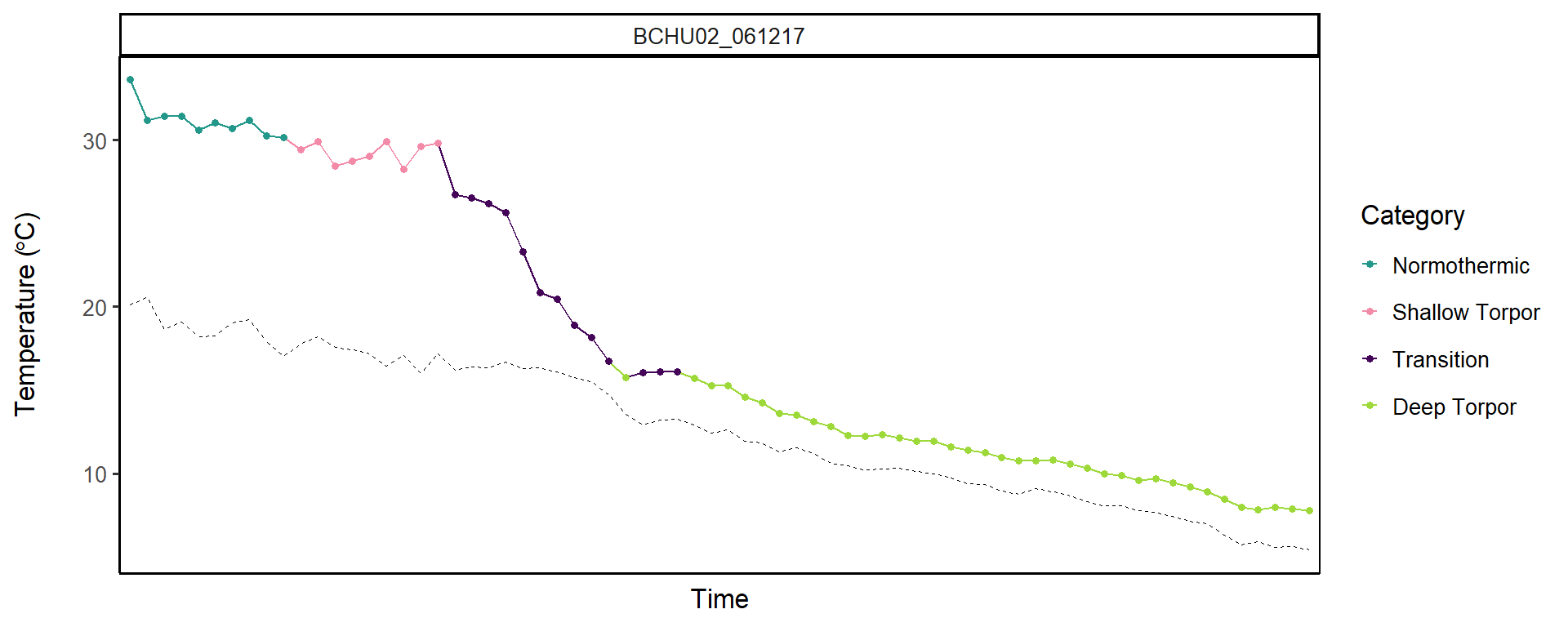

```
##
## [[3]]
```

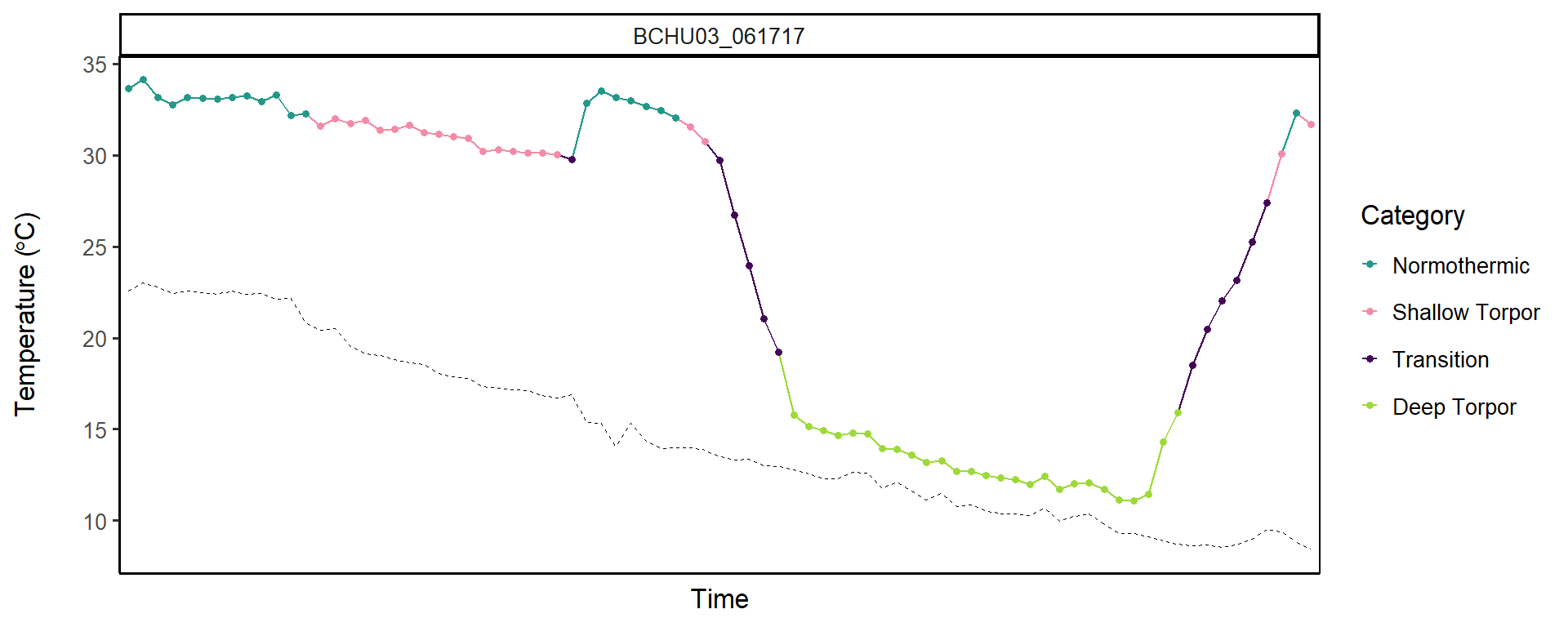

```
##
## [[4]]
```

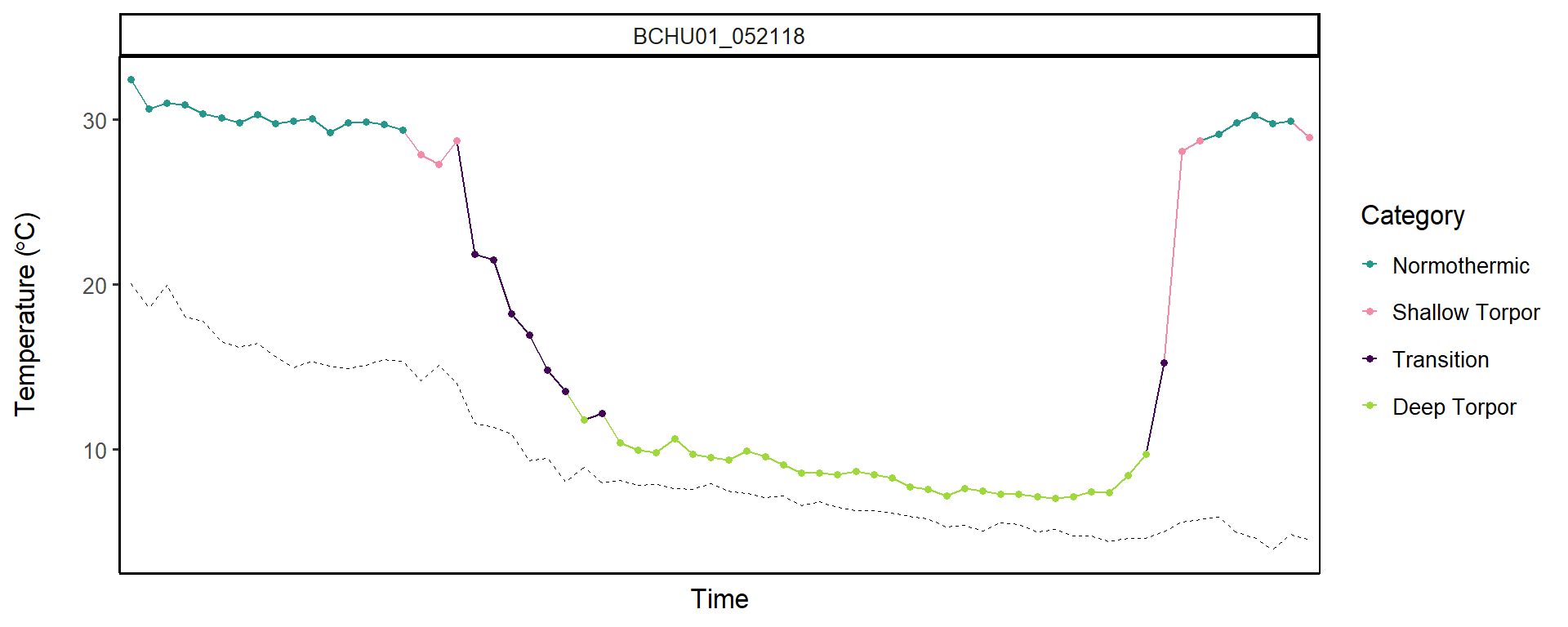

```
##
## [[5]]
```

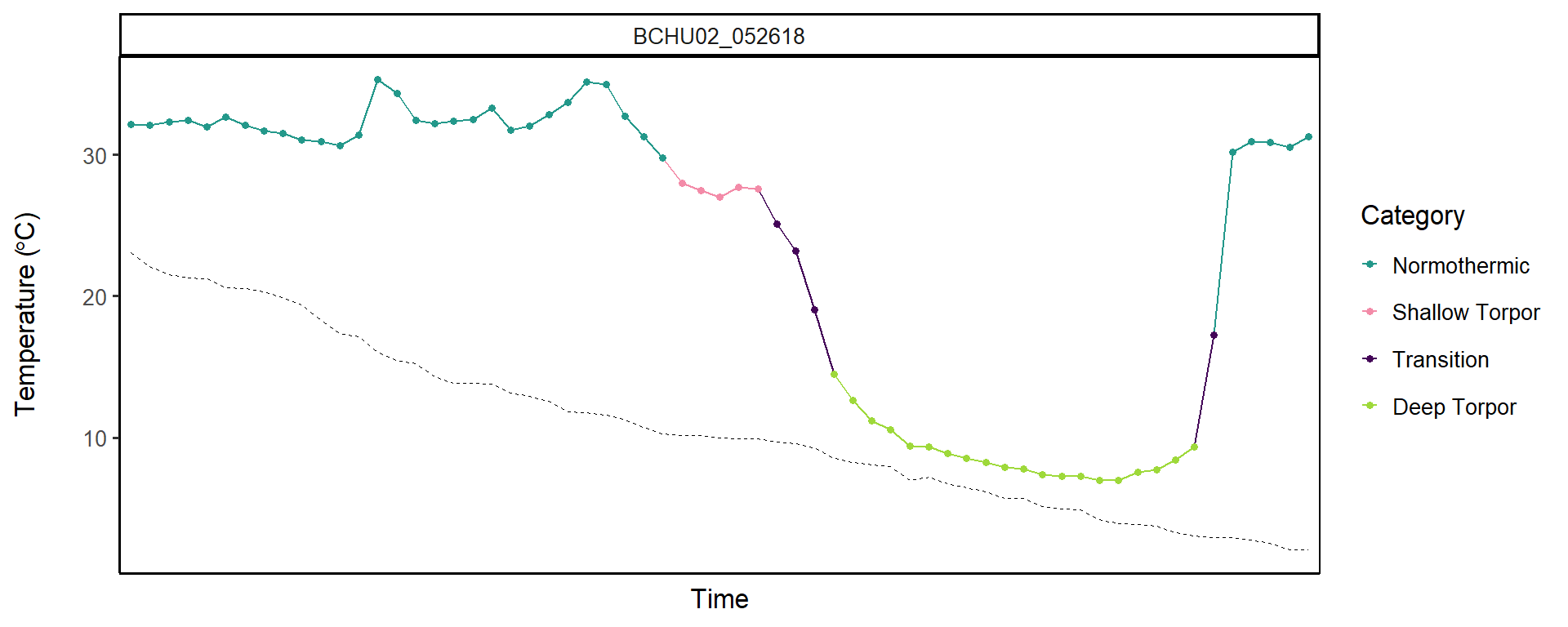

```
##
## [[6]]
```

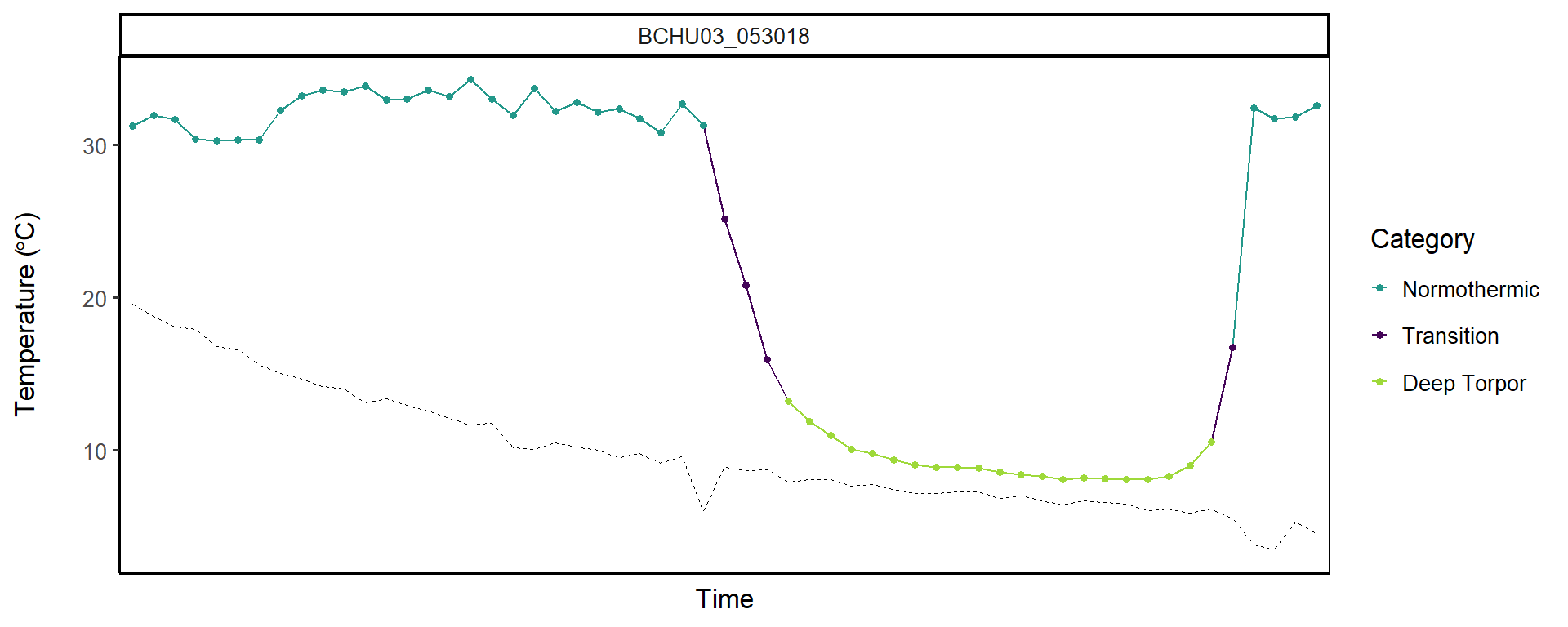

```
##
## [[7]]
```

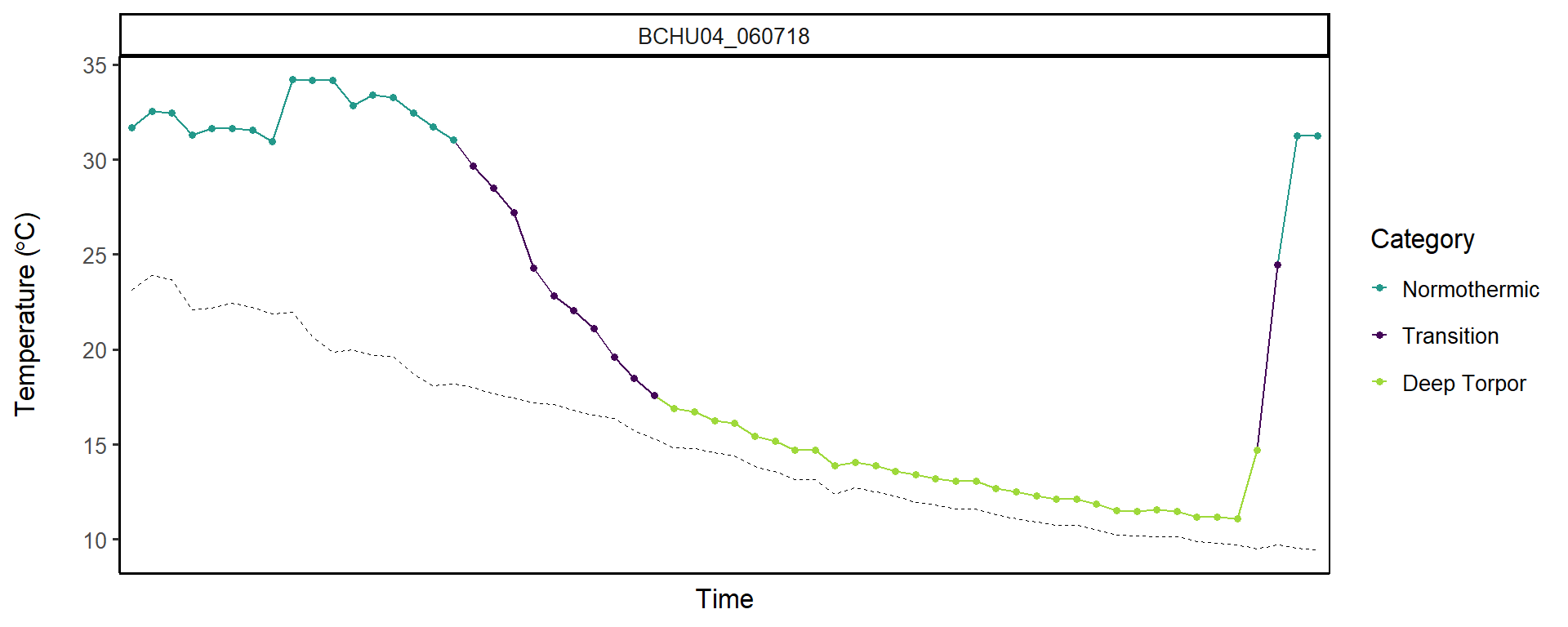

```
##
## [[8]]
```

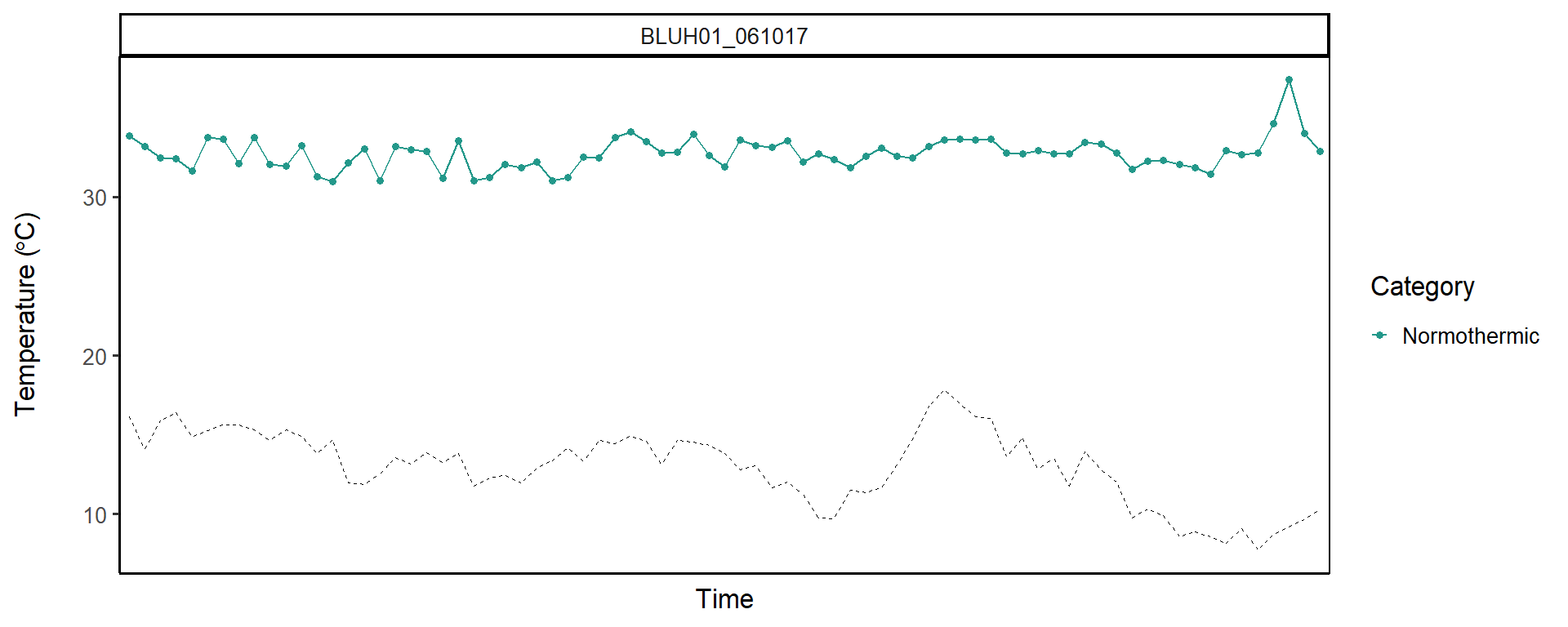

```
##
## [[9]]
```

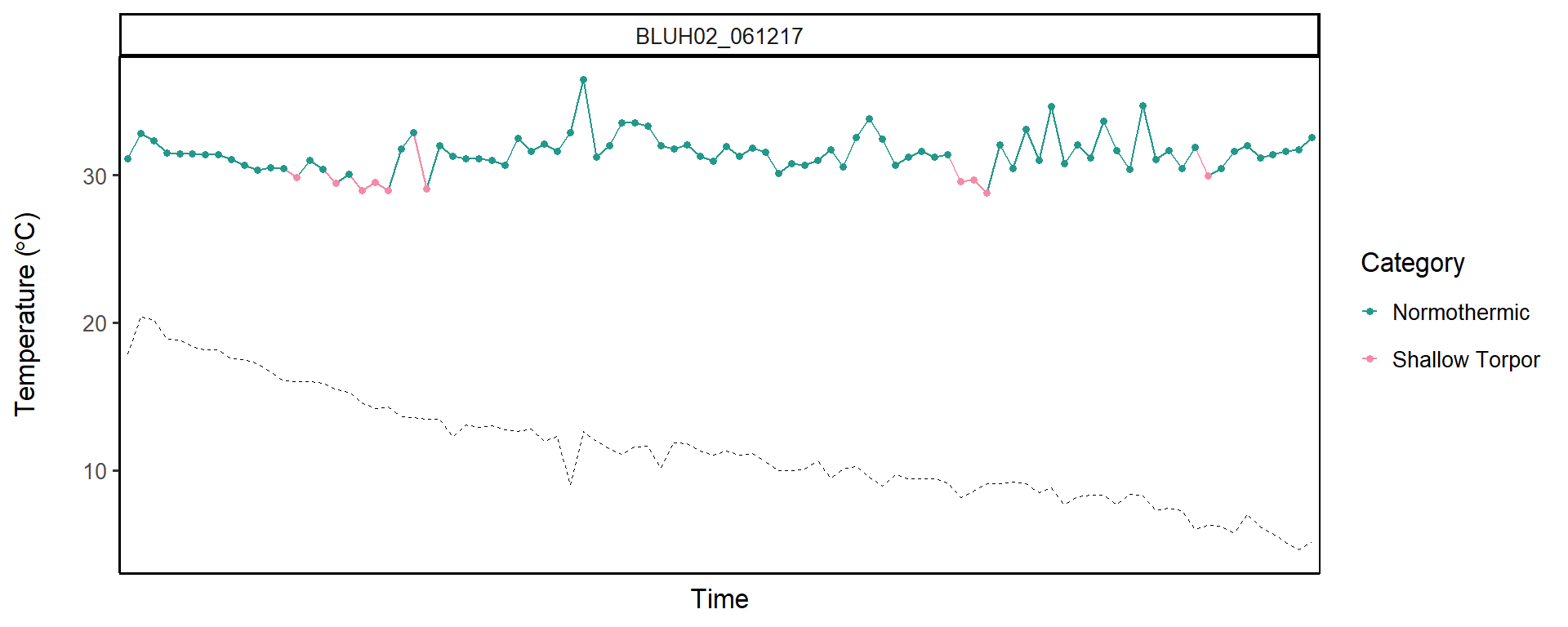

```
##
## [[10]]
```

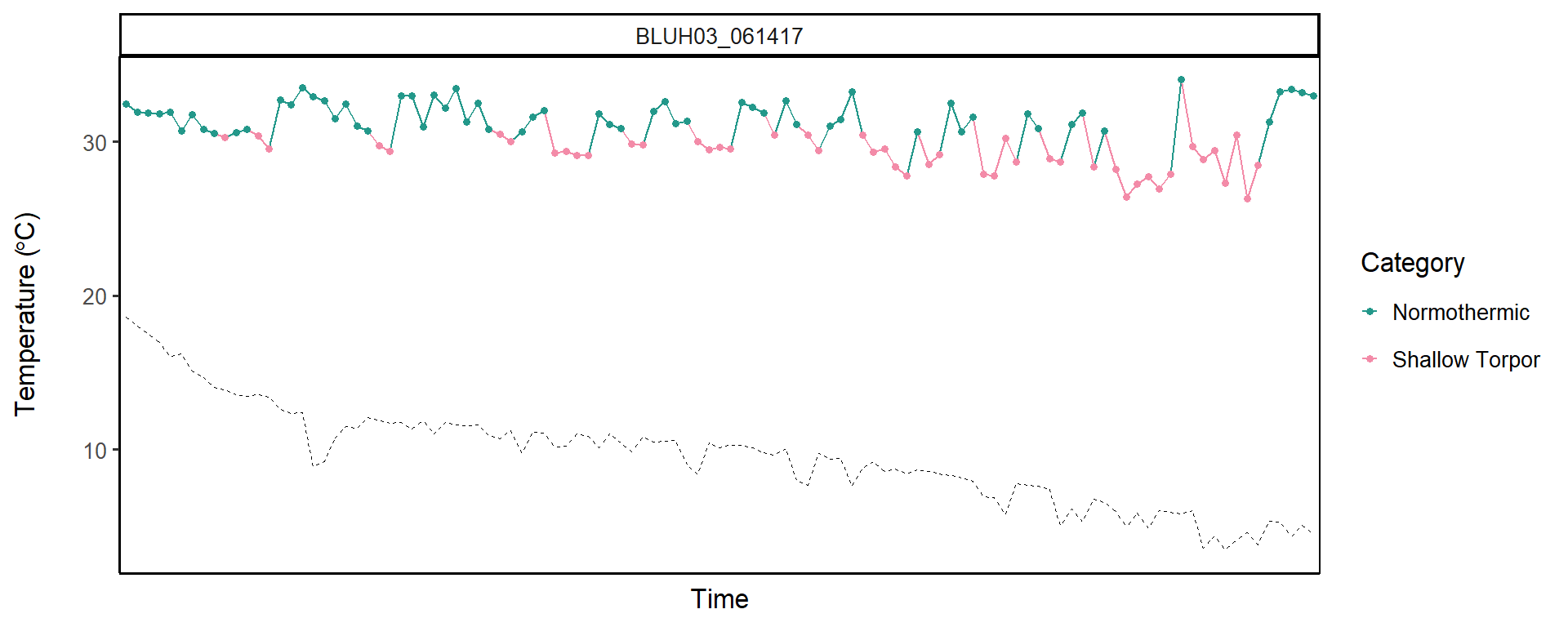

```
##
## [[11]]
```

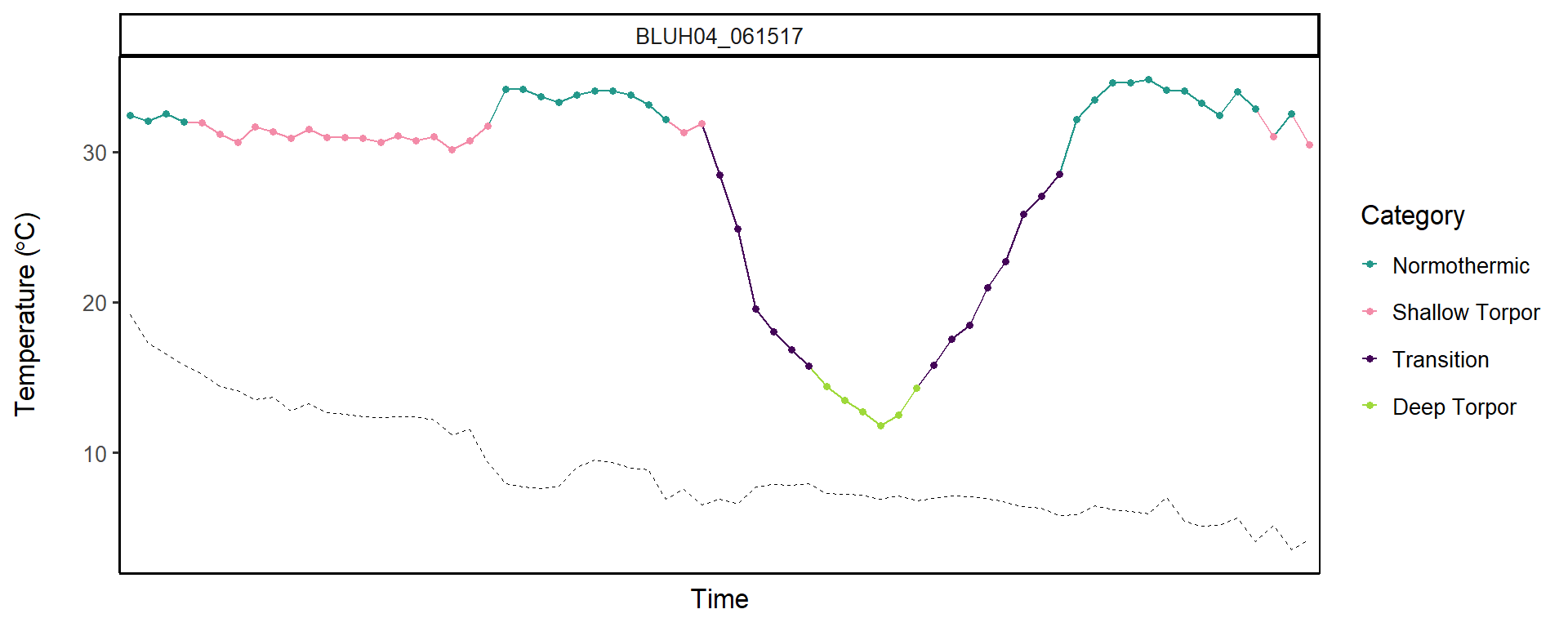

```
##
## [[12]]
```

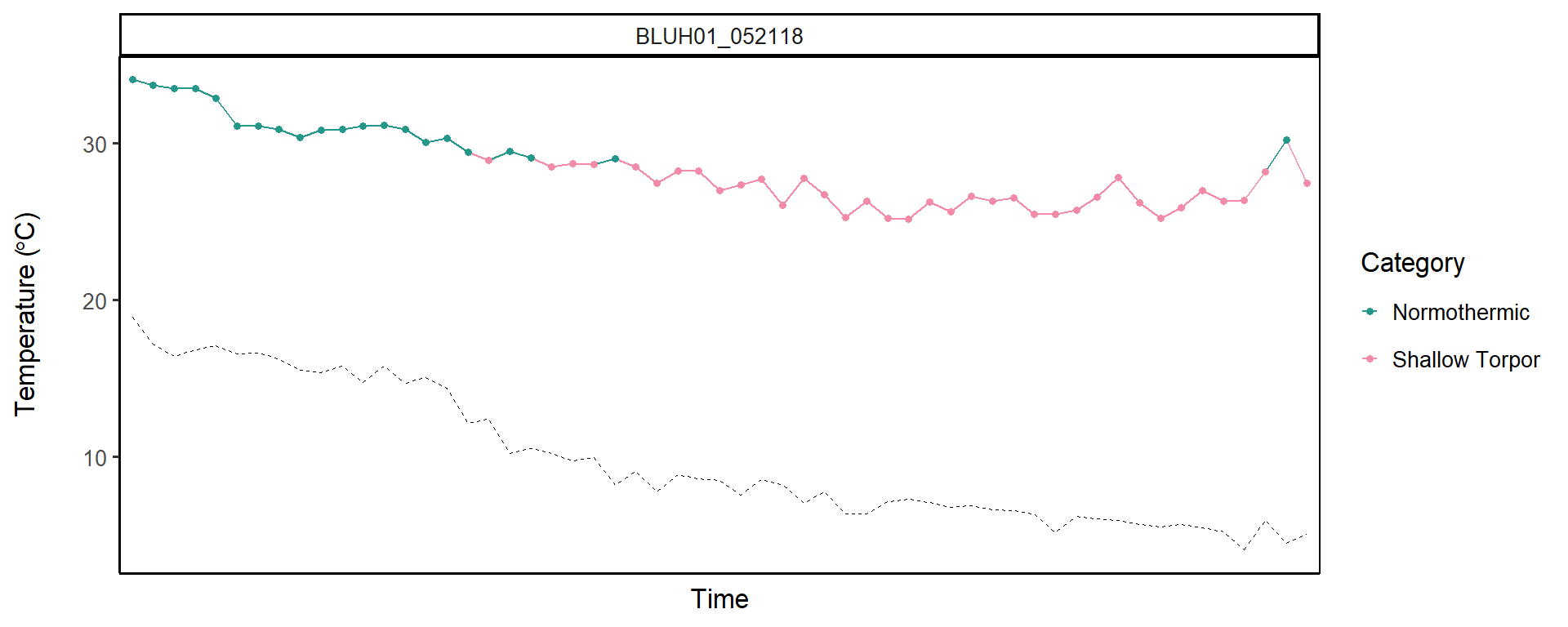

```
##
## [[13]]
```

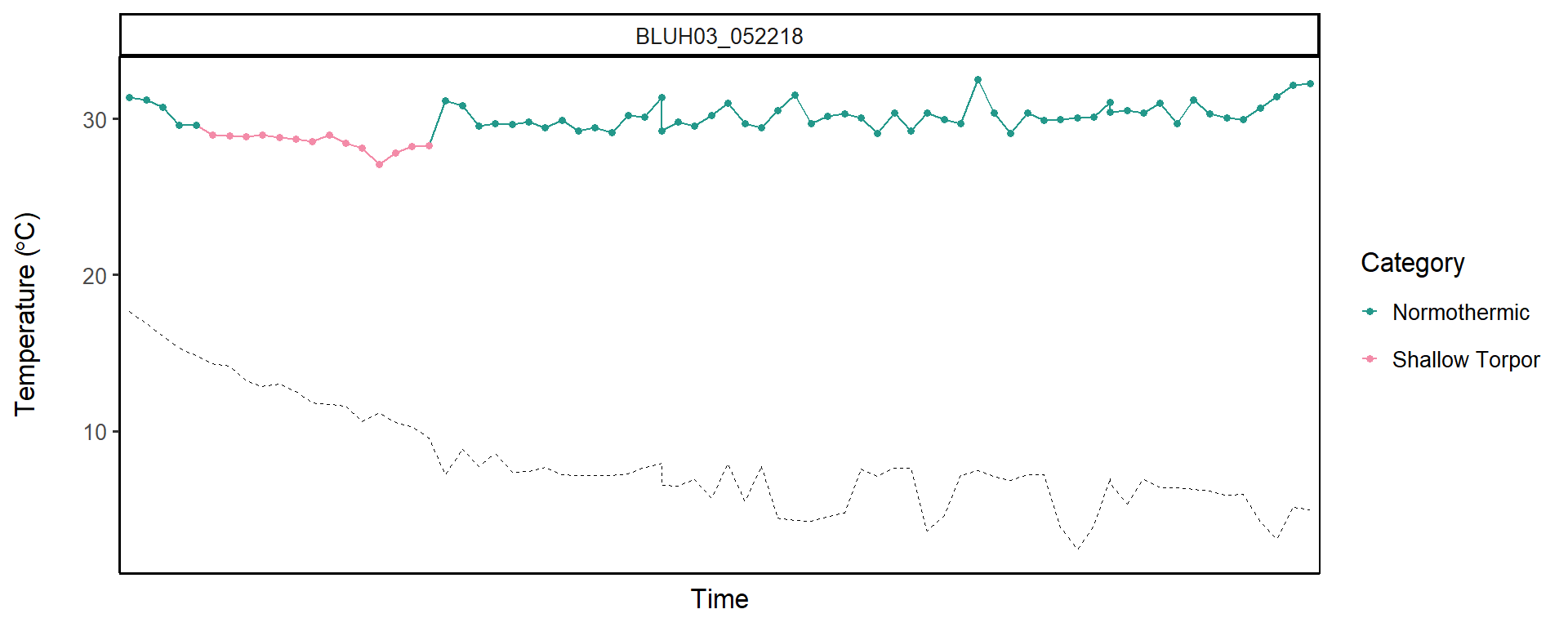

```
##
## [[14]]
```

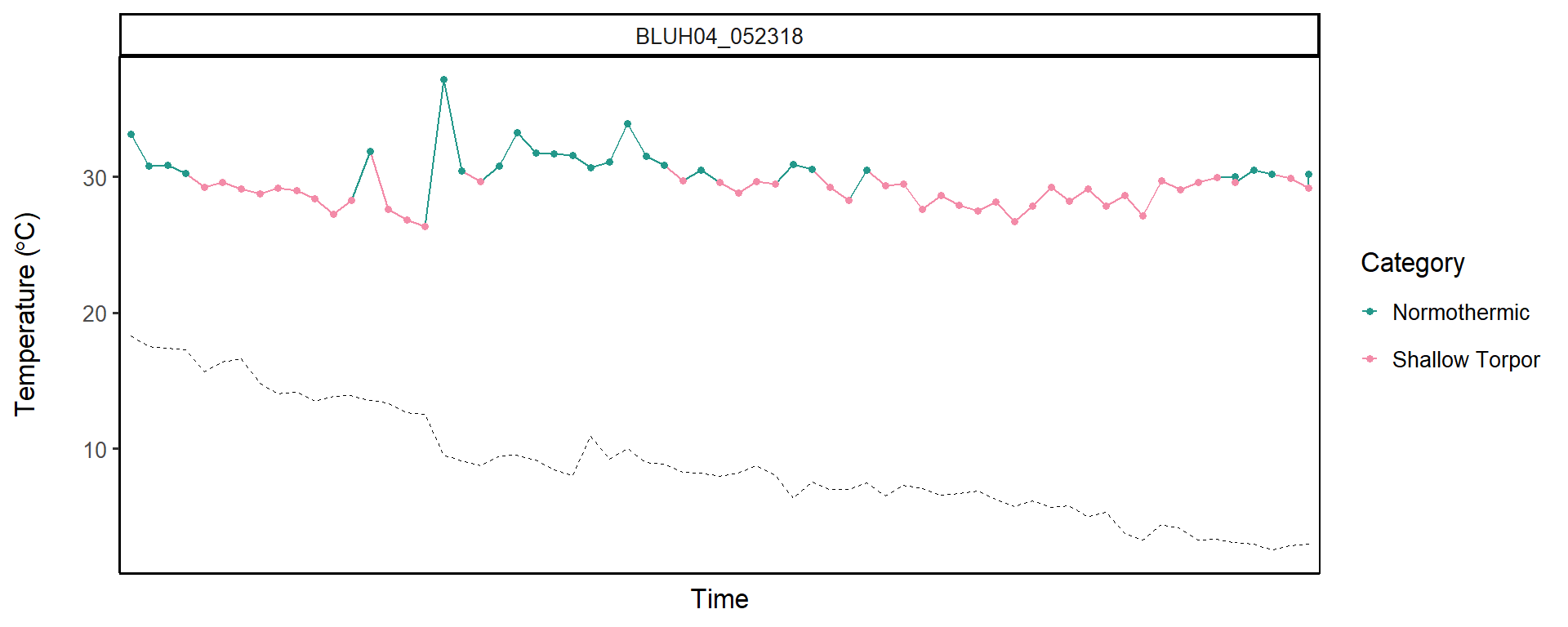

```
##
## [[15]]
```

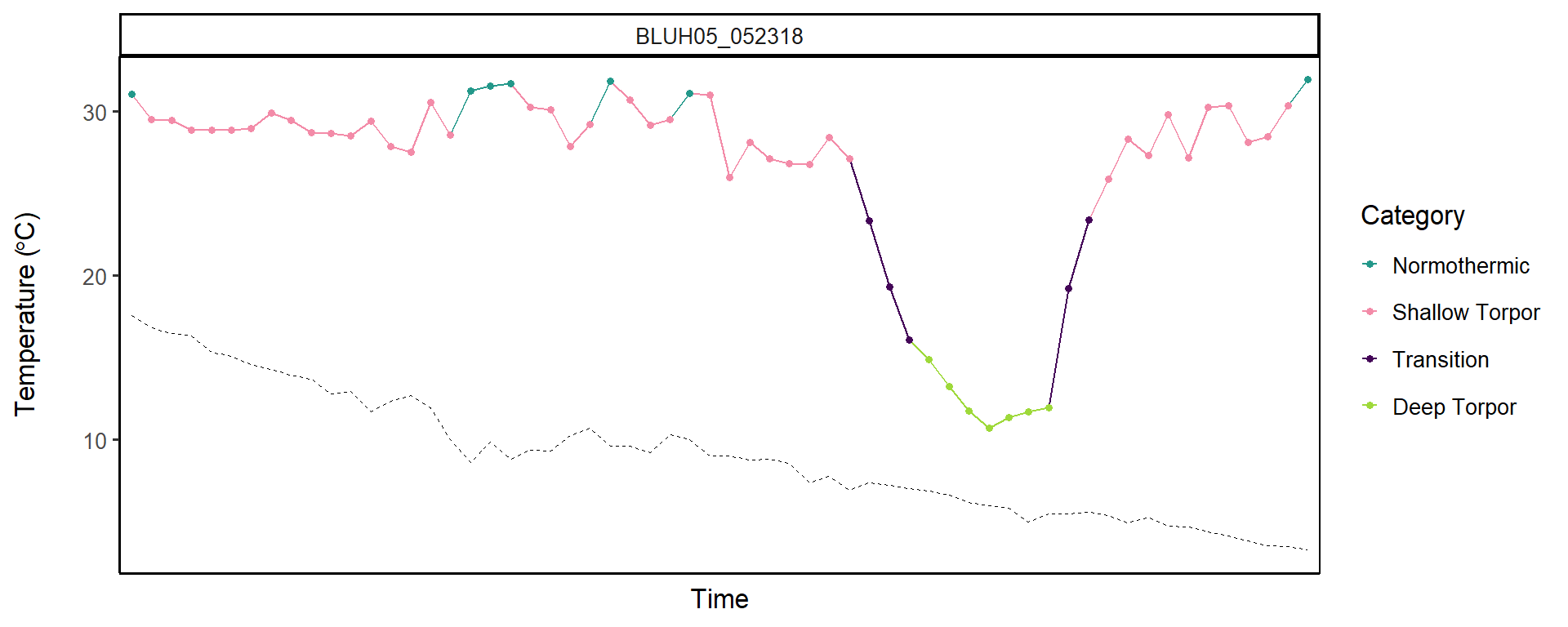

```
##
## [[16]]
```

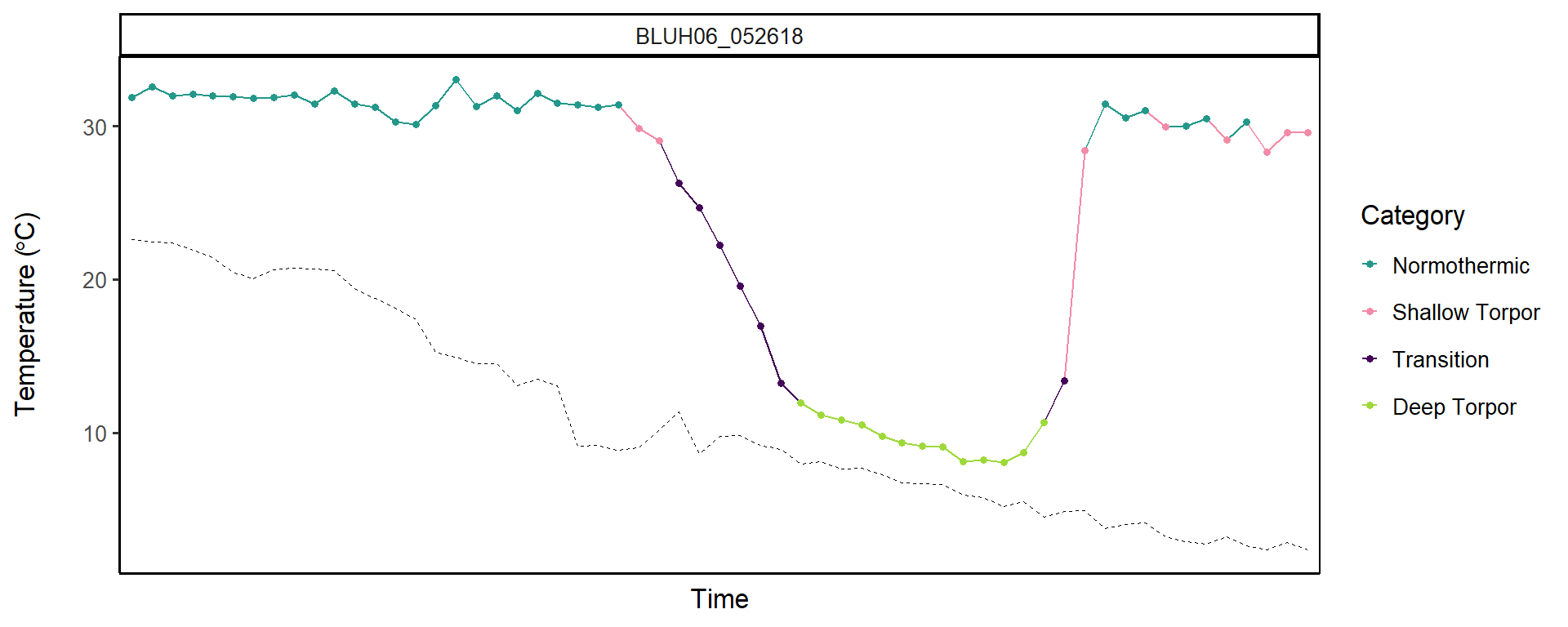

```
##
## [[17]]
```

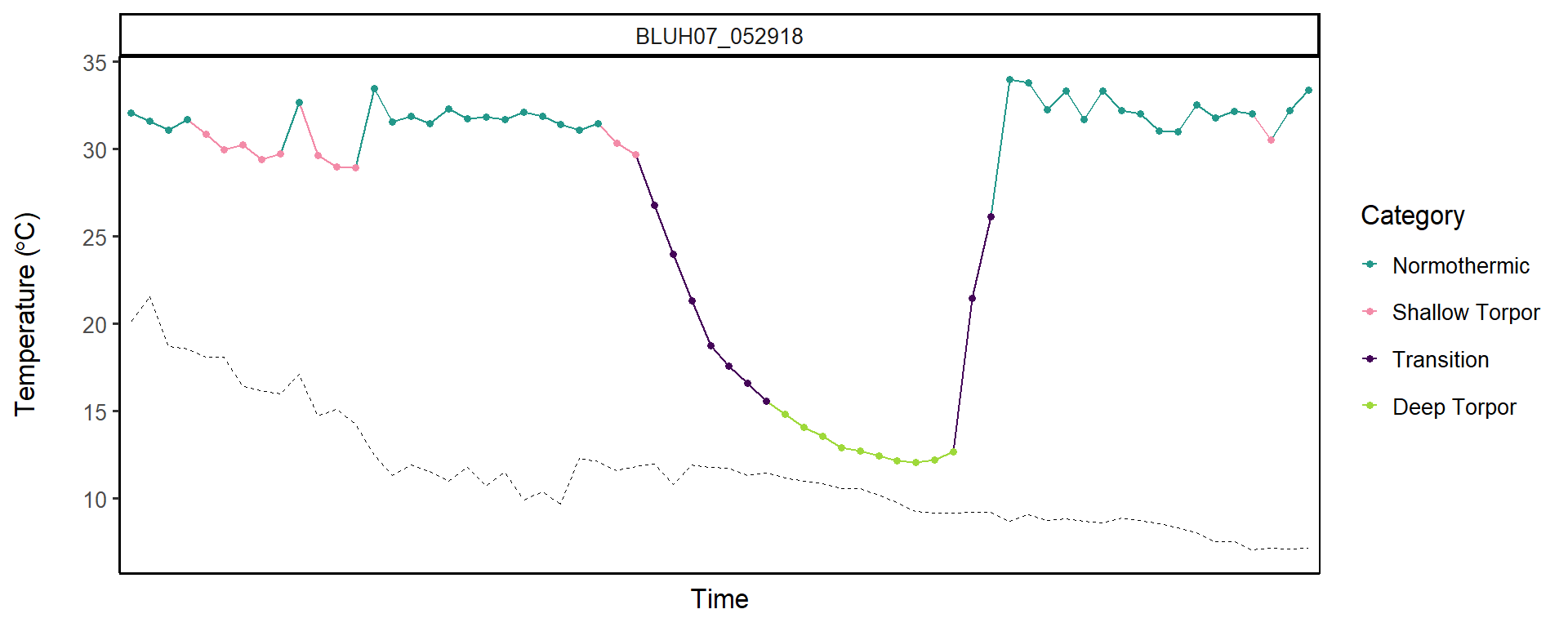

```
##
## [[18]]
```

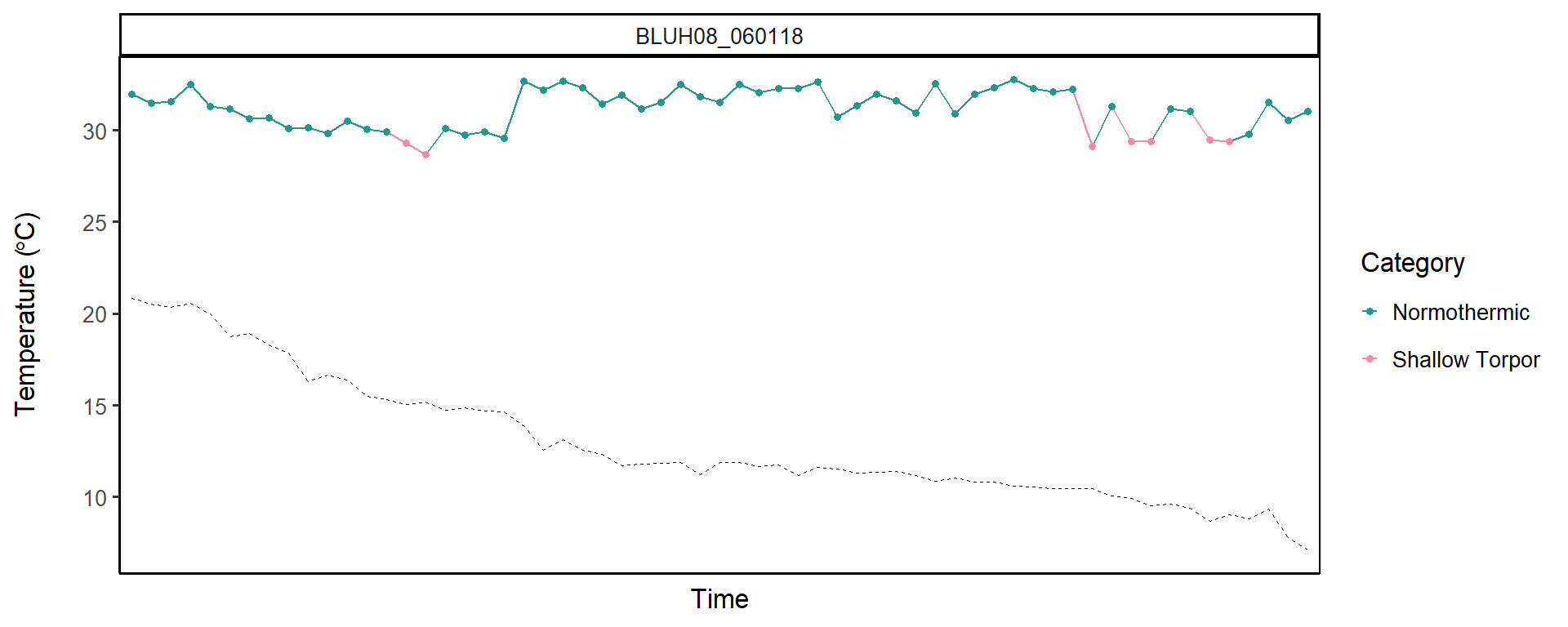

```
##
## [[19]]
```

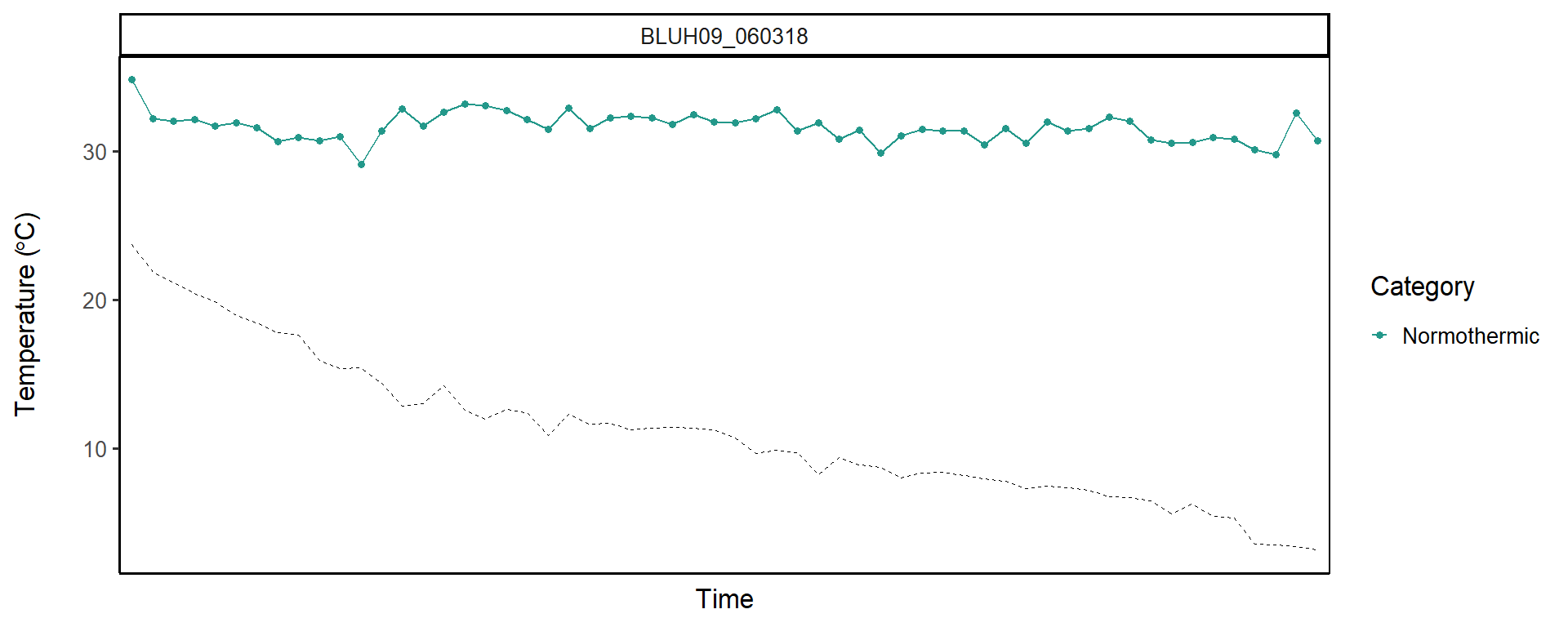

```
##
## [[20]]
```

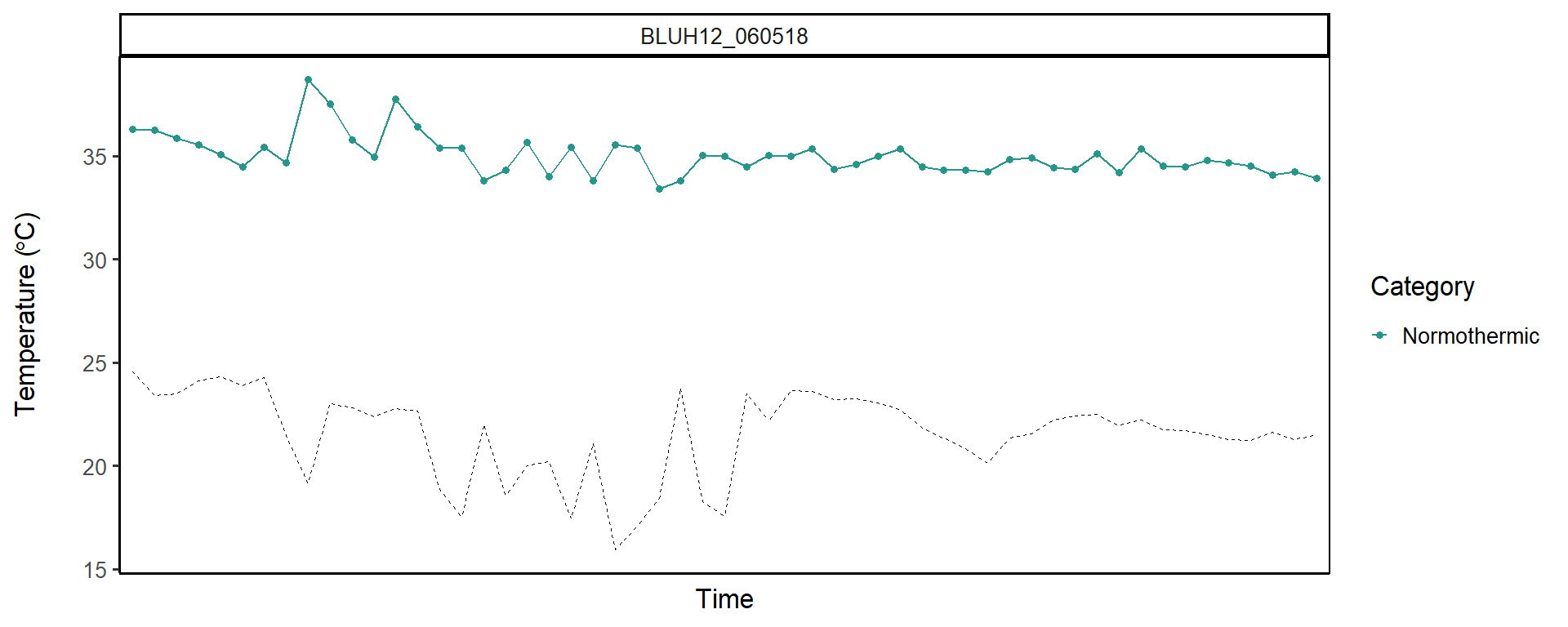

```
##
## [[21]]
```

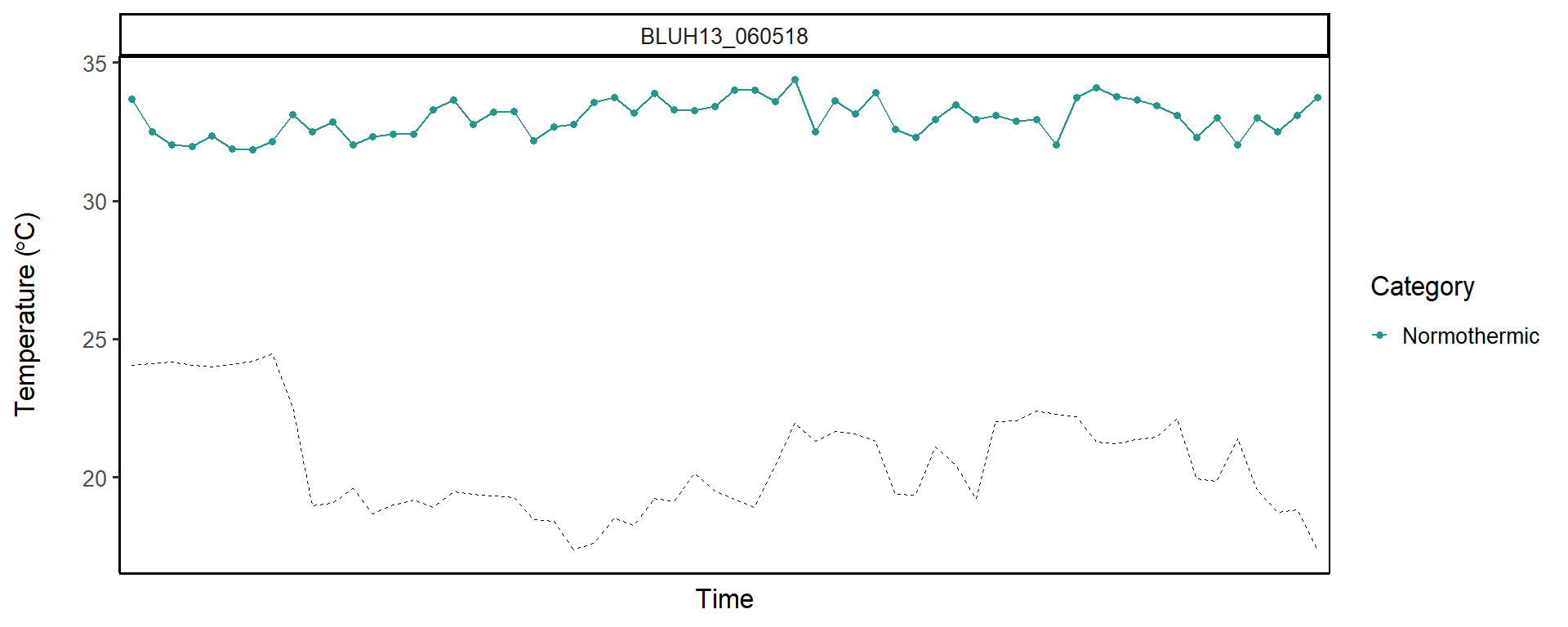

```
##
## [[22]]
```

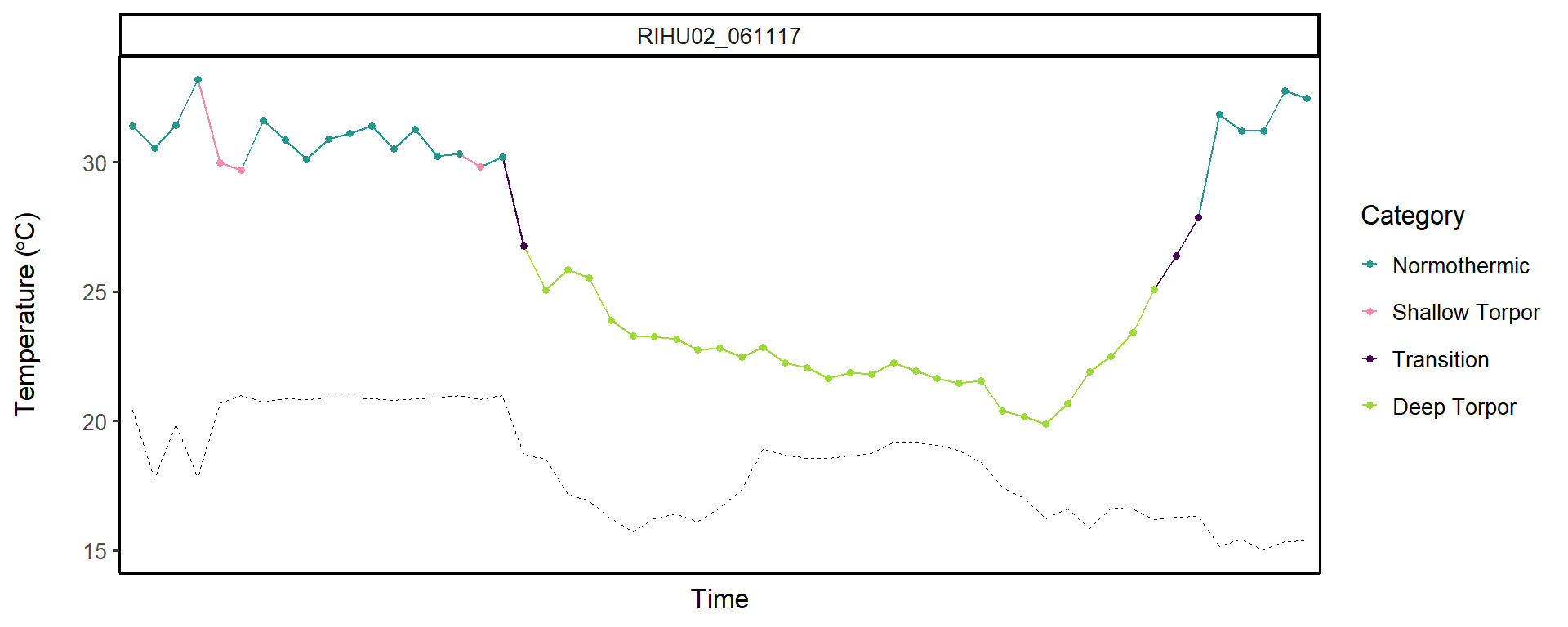

```
##
## [[23]]
```

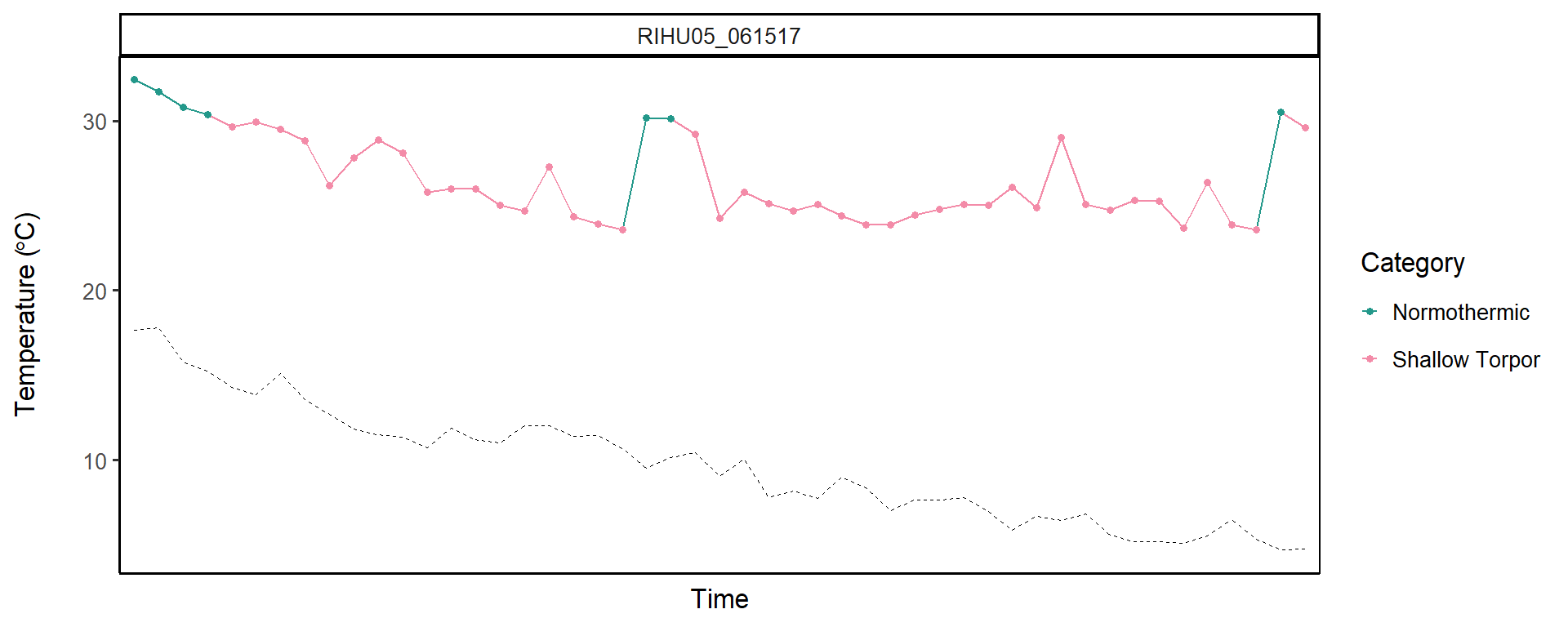

```
##
## [[24]]
```

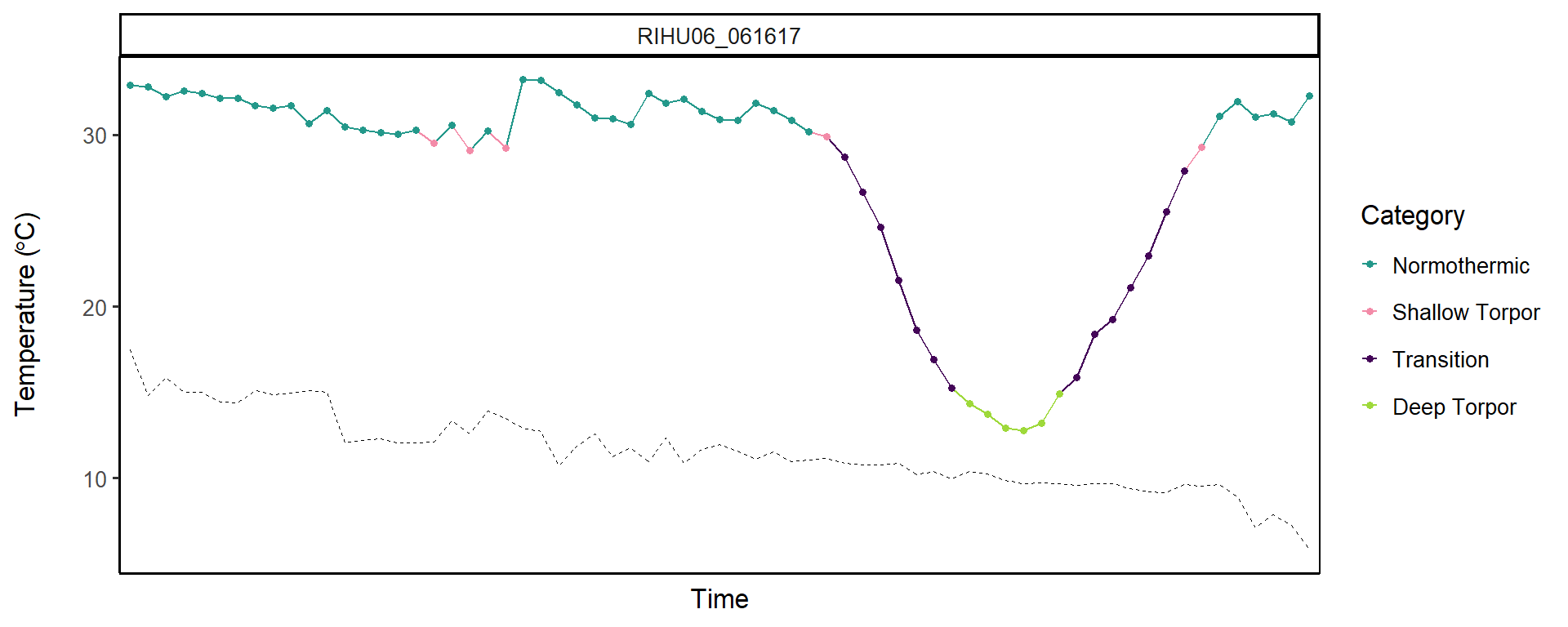

```
##
## [[25]]
```

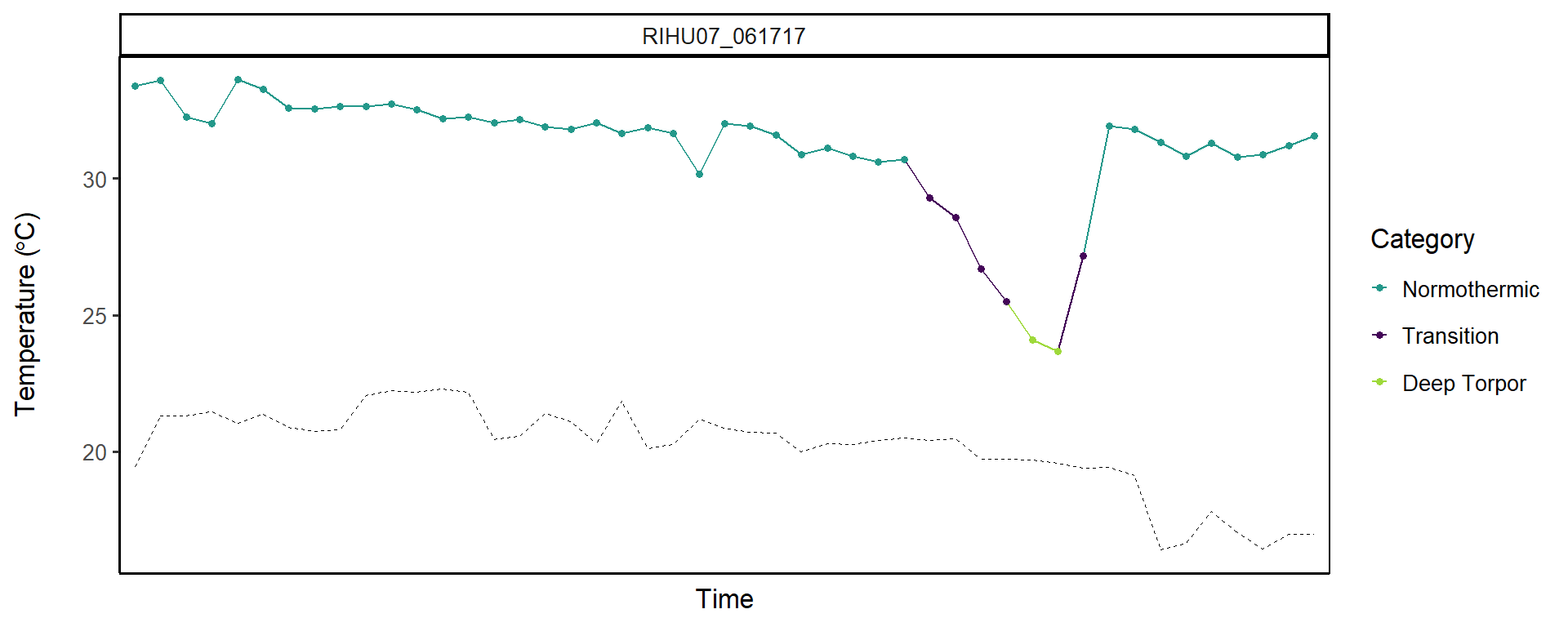

```
##
## [[26]]
```

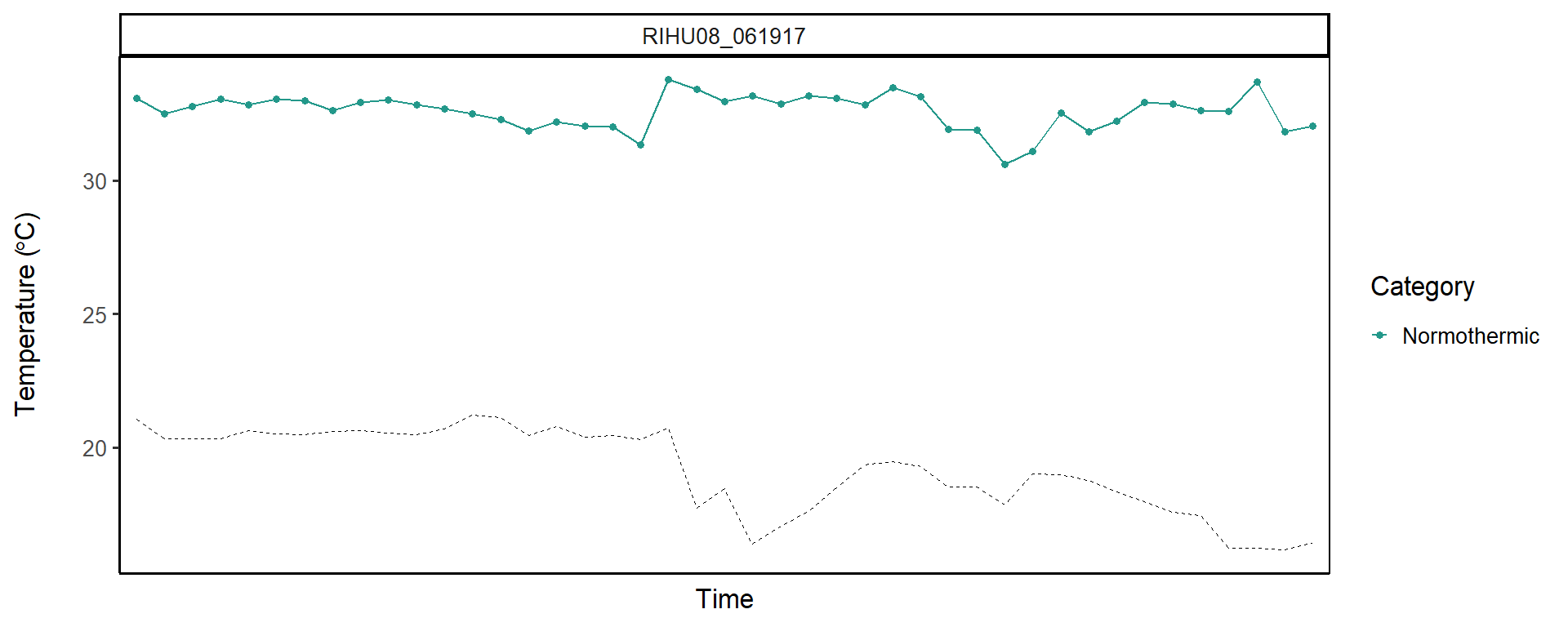

```
##
## [[27]]
```

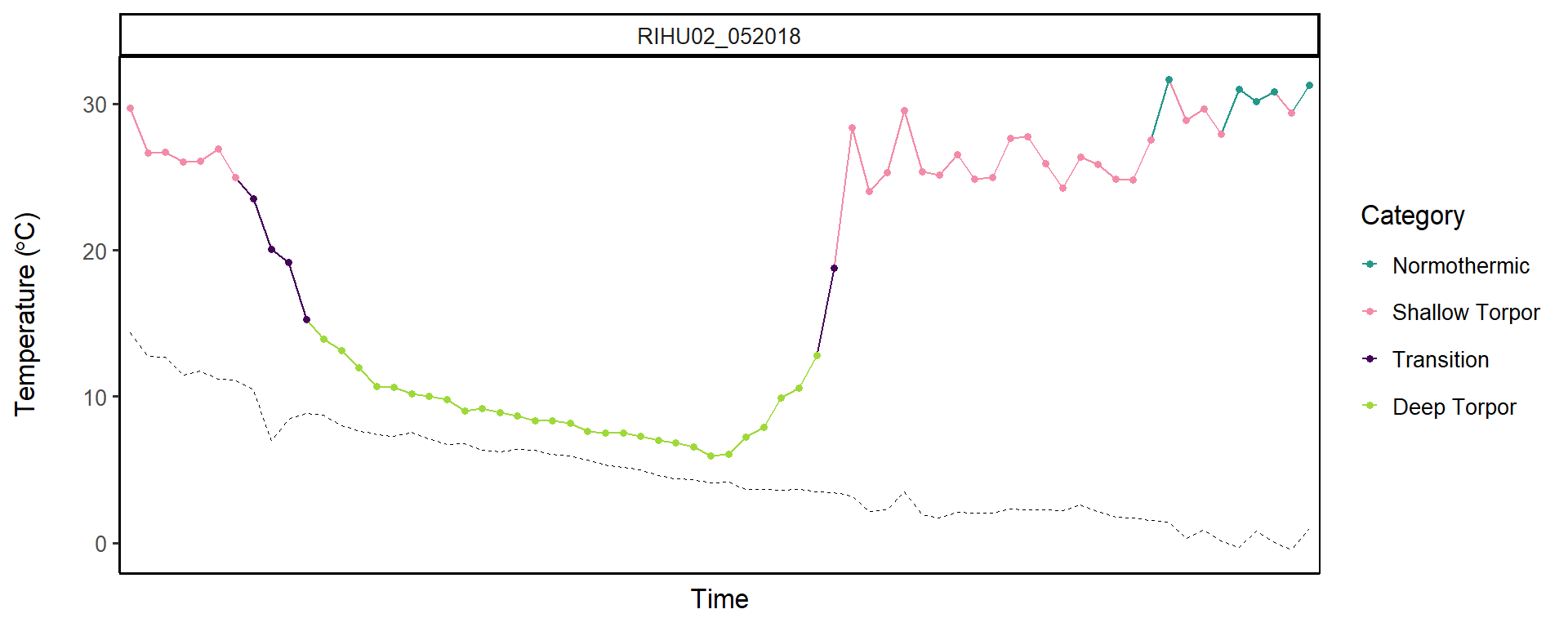

```
##
## [[28]]
```

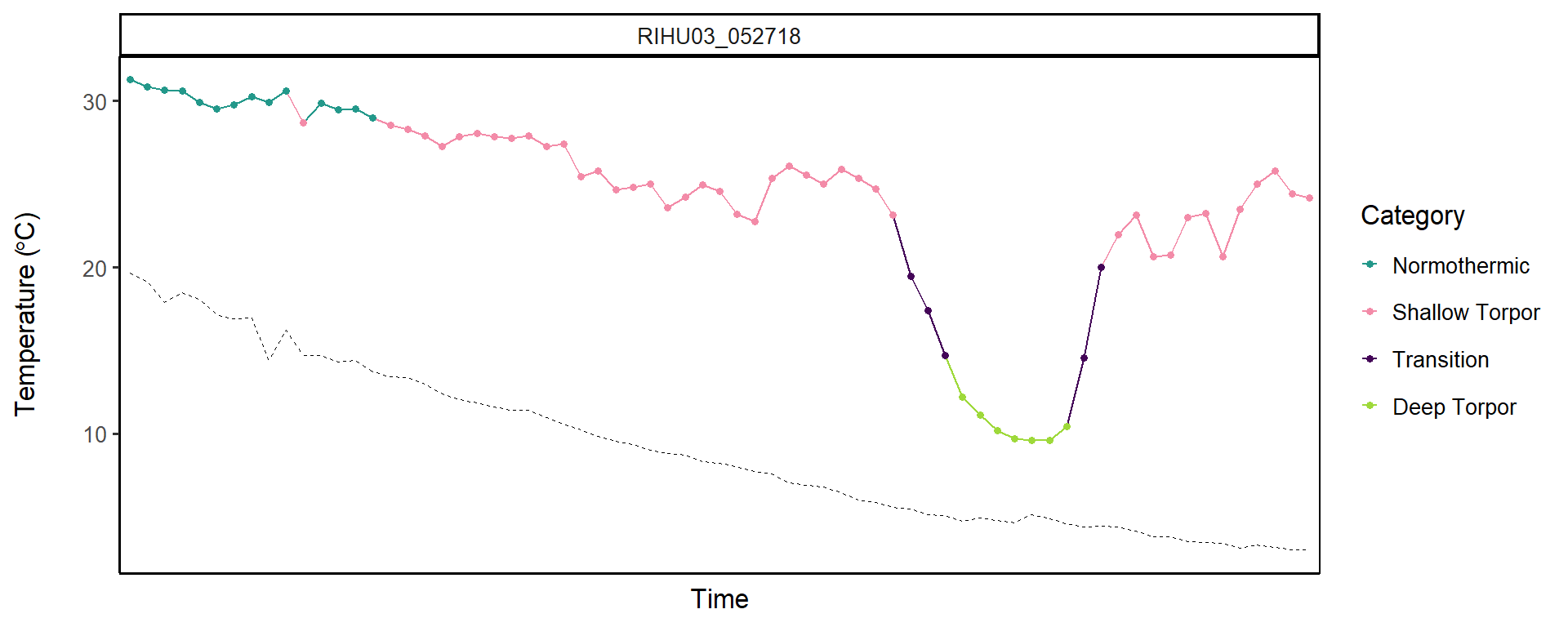

```
##
## [[29]]
```

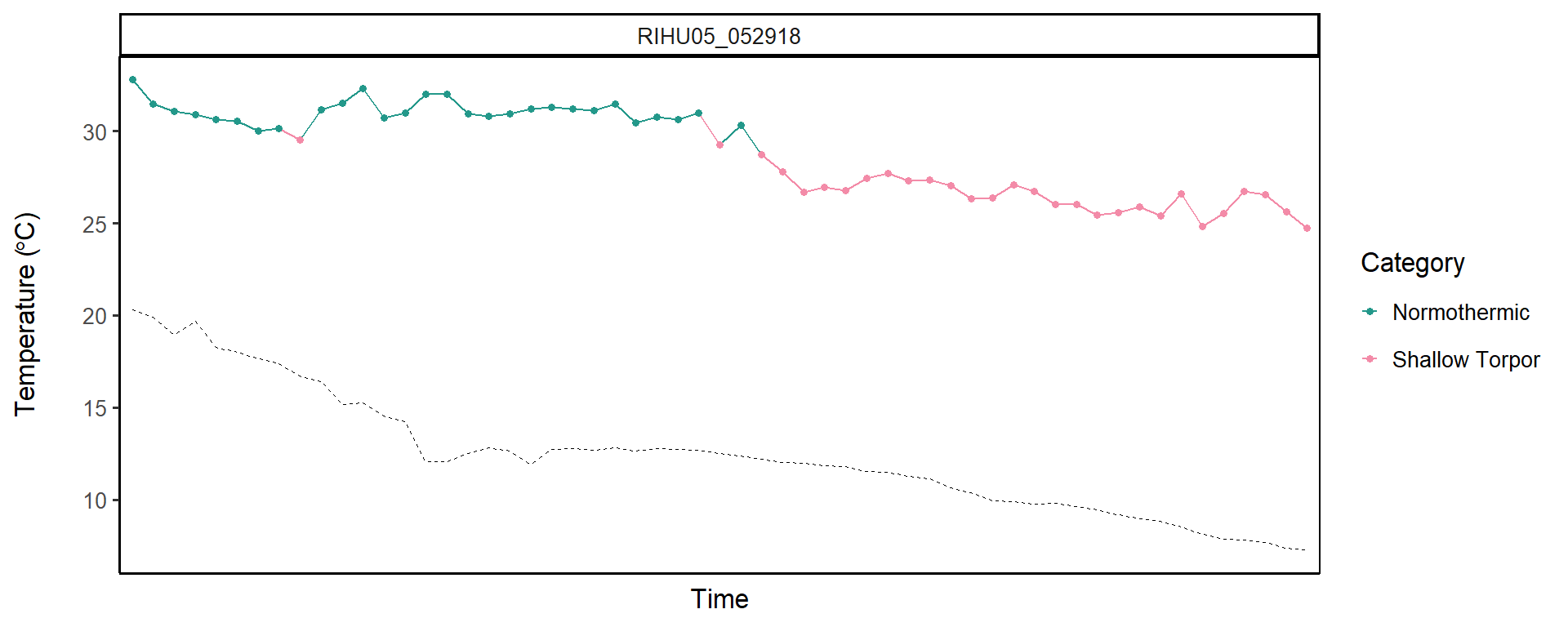

```
##
## [[30]]
```

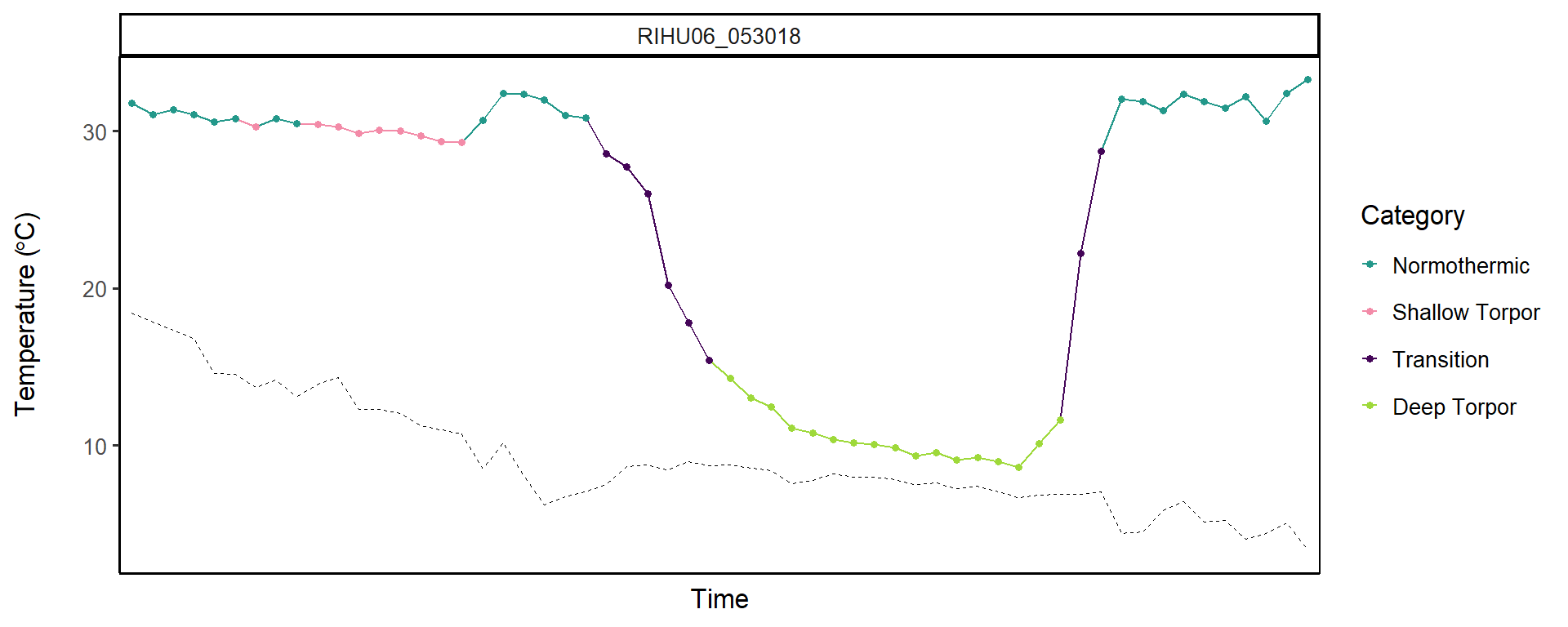

```
##
## [[31]]
```

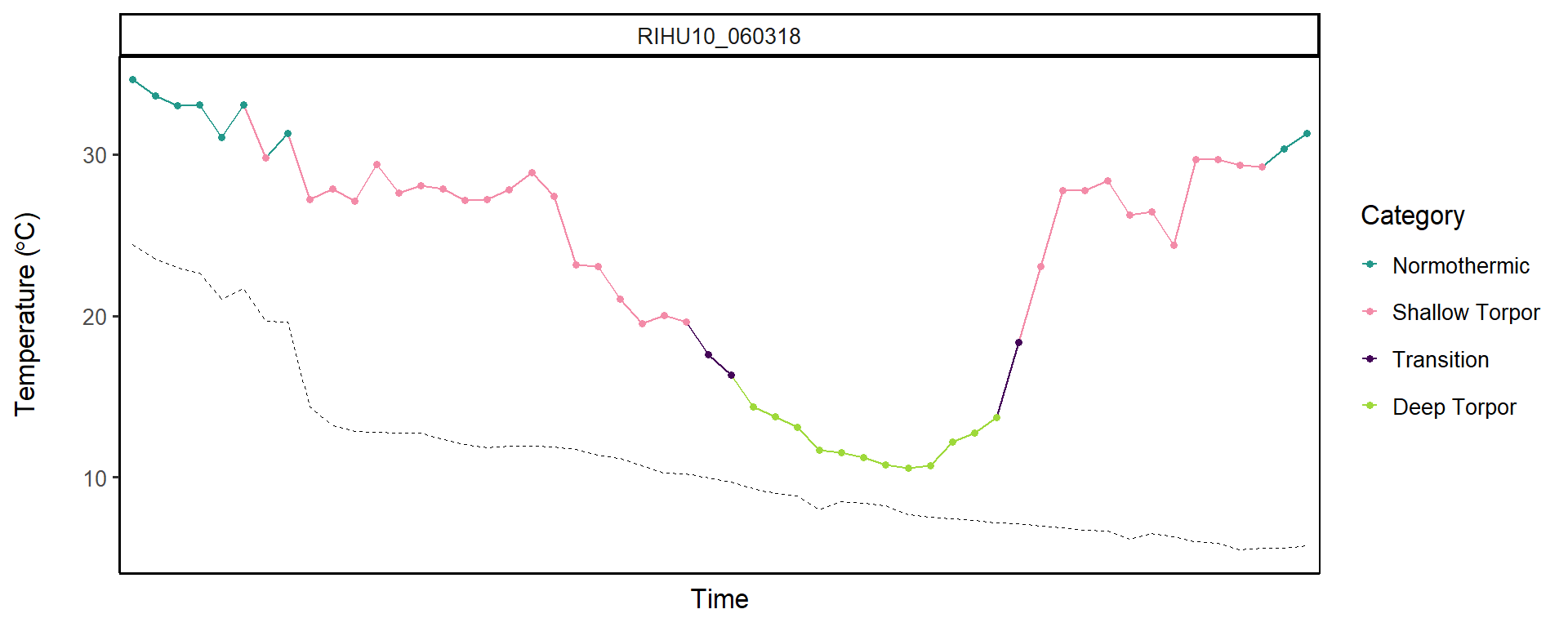

```
##
## [[32]]
```

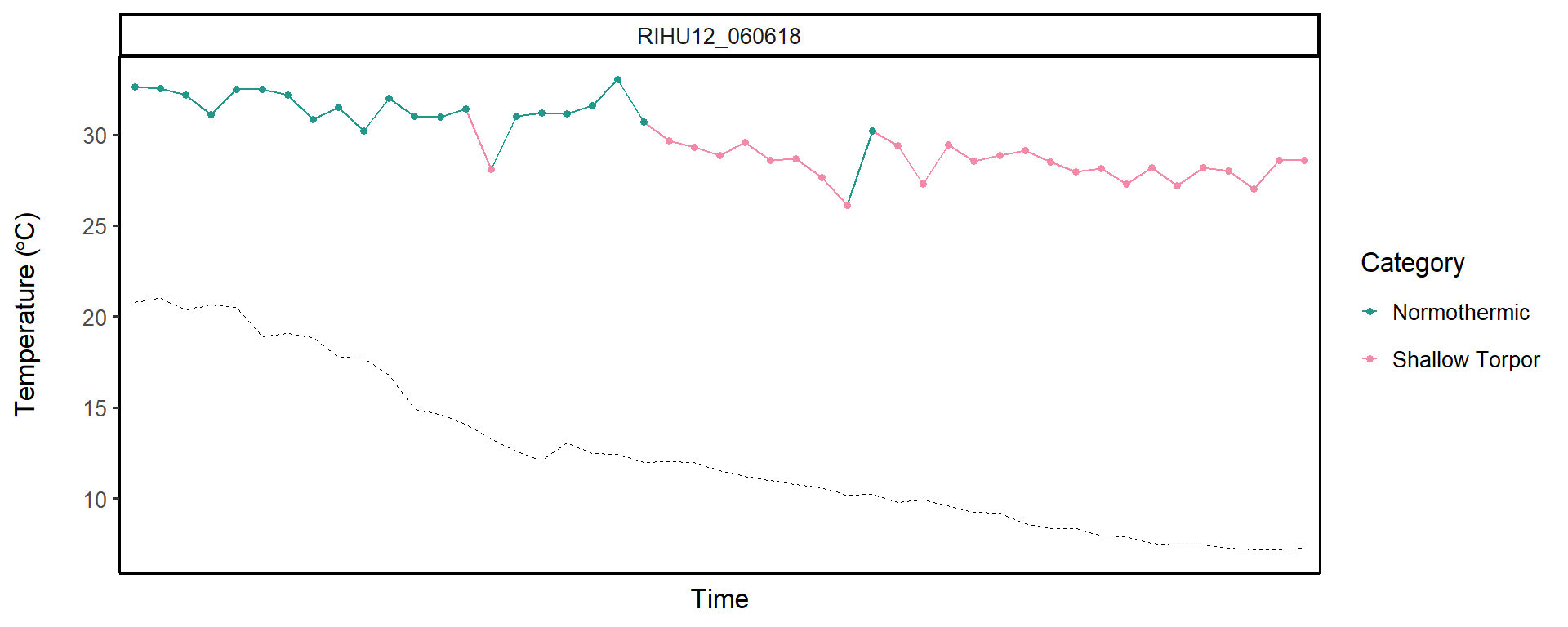

```
##
## [[33]]
```

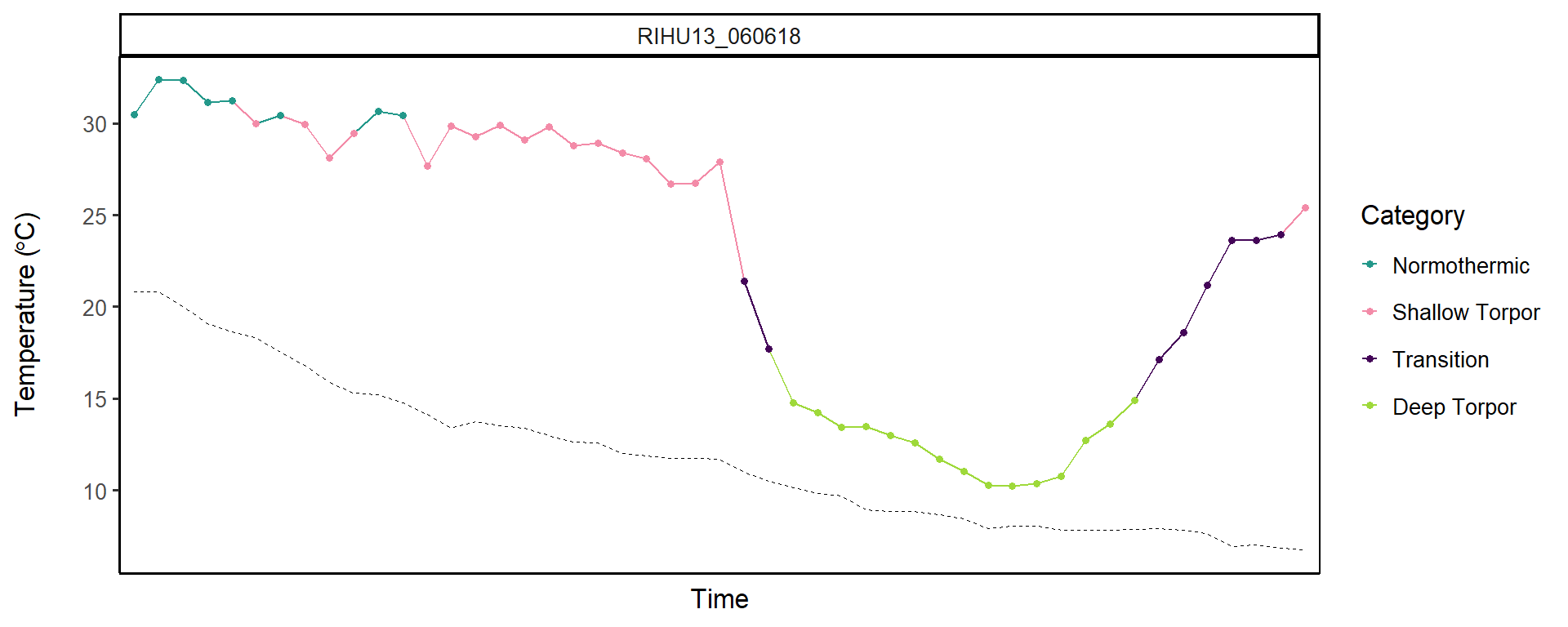
