## Supplement 1 for "Facultative variation across a shallow to deep torpor spectrum in hummingbirds"

Electronic supplementary material for

Supplementary Video SV1: A rotation of a 3D thermal image of a hummingbird to a 2D view of distributions of temperatures over the surface of a hummingbird (accompanies Figure 2).

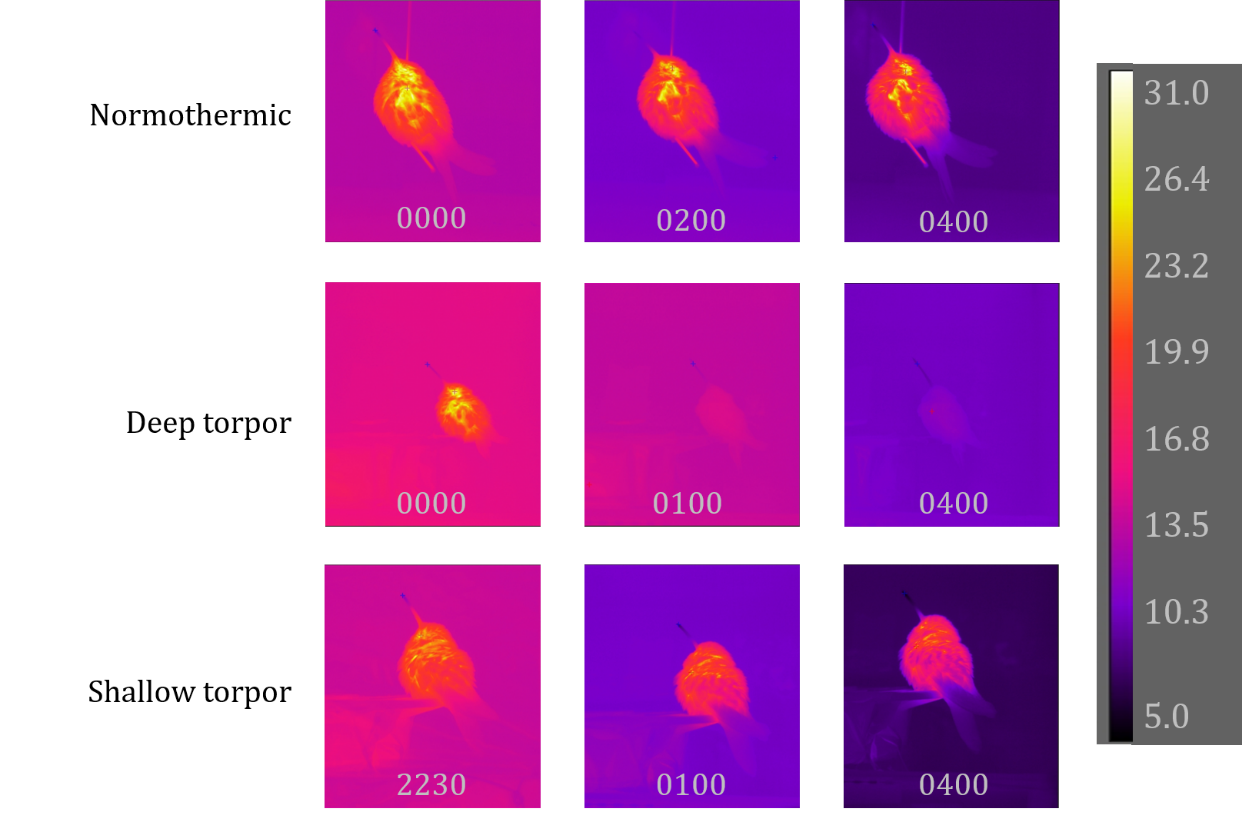

Figure S1: Thermal images of individuals **in all the categories except ‘Transition’. Top: Blue-throated mountain-gem that remained normothermic all night (Surface temperature peaking at 31-32 °C). Middle: black-chinned hummingbird that was normothermic at midnight and then entered deep torpor and was torpid the rest of the night (15 ^°C^). Bottom: Rivoli’s hummingbird that was normothermic at 2230, but then entered shallow torpor for much of the night.**

Models of frequency of use of a thermal category: We first ran a Poisson distribution because the data were a form of frequency or count data (frequency of categories per species across all nights), but the Poisson caused overdispersion, with the residual variance being much higher than the degrees of freedom (see Table S1). We then fit a quasipoisson model, but the model was still overdispersed, with a dispersion parameter of 12.54 (a dispersion parameter great than one means the model is overdispersed). We therefore fit a negative binomial, which resulted in a dispersion parameter of 0.69 (Table S2). This model was therefore a much better fit than either of the others.

Table S1: Model results of the Poisson model **of differences between the species’ use of the four thermal categories (normothermic, shallow, transition, deep torpor). The model was** $Category frequency \sim Category*Species - 1$ **such that intercepts and slopes were allowed to vary by category and by species. The estimates were very similar to those of the negative binomial model, but this model was overdispersed, and was therefore not used.**

| **Category** | **Species** | **Estimate** | **SE** | **Lower CL** | **Upper CL** |
| --- | --- | --- | --- | --- | --- |
| Normothermic | BCHU | 3.51 | 0.07 | 3.39 | 3.64 |
| Shallow Torpor | BCHU | 1.55 | 0.17 | 1.21 | 1.89 |
| Transition | BCHU | 2.52 | 0.11 | 2.31 | 2.73 |
| Deep Torpor | BCHU | 3.90 | 0.05 | 3.79 | 4.00 |
| Normothermic | BLUH | 4.20 | 0.03 | 4.14 | 4.27 |
| Shallow Torpor | BLUH | 3.24 | 0.05 | 3.13 | 3.34 |
| Transition | BLUH | 1.17 | 0.15 | 0.88 | 1.46 |
| Deep Torpor | BLUH | 1.46 | 0.13 | 1.20 | 1.71 |
| Normothermic | RIHU | 3.76 | 0.04 | 3.67 | 3.85 |
| Shallow Torpor | RIHU | 3.48 | 0.05 | 3.38 | 3.58 |
| Transition | RIHU | 2.06 | 0.10 | 1.86 | 2.26 |
| Deep Torpor | RIHU | 2.82 | 0.07 | 2.69 | 2.96 |
| Null deviance: 19037.3 on 132 degrees of freedom  Dispersion parameter for Poisson family taken to be 1 | | | | | |
| Residual deviance: 1905.9 on 120 degrees of freedom  AIC: 2407.7 | | | | | |

Table S2: Model results of the negative binomial model **testing differences between the species’ use of the four thermal categories (normothermic, shallow, transition, deep torpor). The model was** $Category proportion \sim Category*Species - 1$ **such that intercepts and slopes were allowed to vary by category and by species. These frequency estimates are only meaningful in relation to one another, and can be compared by converting them into proportions within a species (as depicted in Figure 5b on the right). All the estimates’ confidence intervals do not overlap with zero.**

| **Species** | **Estimate** | **SE** | **Lower CL** | **Upper CL** |
| --- | --- | --- | --- | --- |
| BCHU Normothermic | 3.51 | 0.47 | 2.59 | 4.43 |
| BCHU Shallow Torpor | 1.55 | 0.50 | 0.58 | 2.52 |
| BCHU Transition | 2.52 | 0.48 | 1.59 | 3.45 |
| BCHU Deep Torpor | 3.90 | 0.47 | 2.98 | 4.81 |
| BLUH Normothermic | 4.20 | 0.33 | 3.56 | 4.85 |
| BLUH Shallow Torpor | 3.24 | 0.33 | 2.58 | 3.89 |
| BLUH Transition | 1.17 | 0.36 | 0.46 | 1.87 |
| BLUH Deep Torpor | 1.46 | 0.35 | 0.76 | 2.15 |
| RIHU Normothermic | 3.76 | 0.36 | 3.06 | 4.46 |
| RIHU Shallow Torpor | 3.48 | 0.36 | 2.78 | 4.18 |
| RIHU Transition | 2.06 | 0.37 | 1.33 | 2.78 |
| RIHU Deep Torpor | 2.82 | 0.36 | 2.11 | 3.53 |
| Null deviance: 2935.95 on 132 degrees of freedom  Dispersion parameter for Negative Binomial (0.6627) family taken to be 1 | | | | |
| Residual deviance: 157.93 on 120 degrees of freedom  AIC: 1028.6 | | | | |

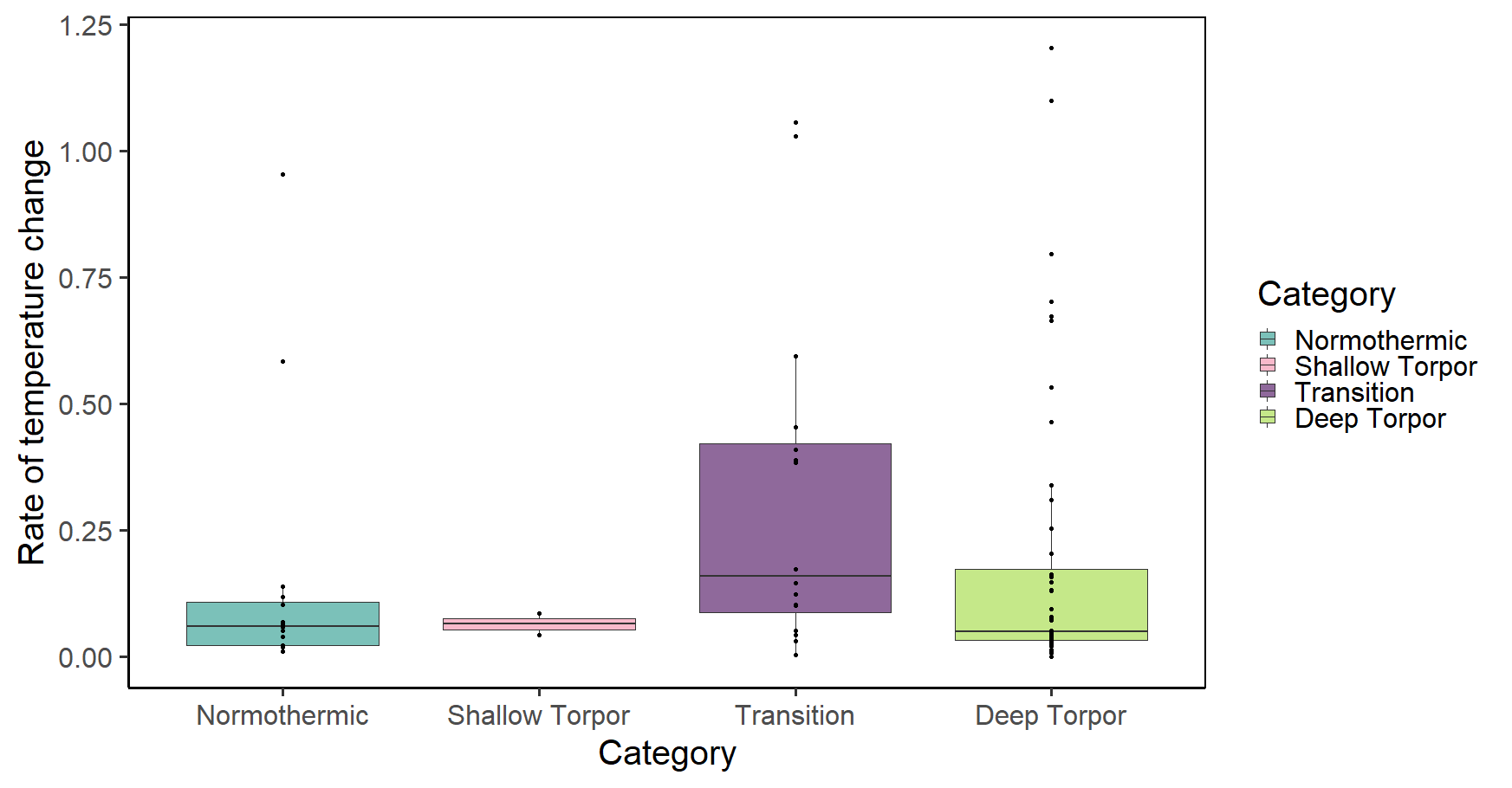

**Figure S2:** Absolute rates of temperature change per category in °C/minute.

Models of surface temperature**:** To first test the effect of ambient temperature on surface temperature, we ran a simple linear model of surface temperature ($T_{S}$) as a function of ambient temperature ($T_{a}$), without incorporating the other variables. This model only explained 15% of the variation in surface temperatures, and we therefore ran a linear mixed effects model of surface temperature as a function of ambient temperature. We included mass as a continuous fixed covariate, and thermal category (normothermy, shallow torpor, etc.), year (because ambient temperatures varied across the two years), and species as discrete covariates. We modelled categories within individuals as a random covariate. We included a correlation structure (‘corAR1’) to account for temporal autocorrelation, with categories nested within individuals; and interaction effects between category and species and between ambient temperature and category.

$$T_{s} \sim T_{a}*Category+ Species*Category+Capture mass+Year+\left( 1 | \frac{Individual}{Category} \right)+CorAR1$$

We tested for normality of the residuals, homogeneity of variances, and linearity to confirm that the model was a good fit.

**Table S3:** Model outputs of the surface temperature model presented above. These are mean estimates of each thermal category per species, with standard error, degrees of freedom, and lower and upper confidence limits.

| **Species** | **Category** | **Mean** | **SE** | **df** | **Lower CL** | **Upper CL** |
| --- | --- | --- | --- | --- | --- | --- |
| BCHU | Normothermic | 33.03 | 1.83 | 28 | 29.29 | 36.77 |
| BLUH | Normothermic | 31.81 | 0.68 | 28 | 30.42 | 33.19 |
| RIHU | Normothermic | 30.93 | 0.52 | 28 | 29.85 | 32.00 |
| BCHU | Shallow Torpor | 30.40 | 1.91 | 28 | 26.48 | 34.32 |
| BLUH | Shallow Torpor | 29.03 | 0.78 | 28 | 27.44 | 30.62 |
| RIHU | Shallow Torpor | 27.48 | 0.51 | 28 | 26.44 | 28.53 |
| BCHU | Transition | 21.56 | 1.85 | 28 | 17.77 | 25.36 |
| BLUH | Transition | 22.07 | 1.00 | 28 | 20.02 | 24.13 |
| RIHU | Transition | 22.71 | 0.66 | 28 | 21.37 | 24.06 |
| BCHU | Deep Torpor | 15.10 | 1.82 | 28 | 11.37 | 18.84 |
| BLUH | Deep Torpor | 15.46 | 1.02 | 28 | 13.38 | 17.54 |
| RIHU | Deep Torpor | 15.67 | 0.59 | 28 | 14.46 | 16.89 |

Table S4: Summary of the full surface temperature model results **for each term in the model. T_a_ = ambient temperature.**

| **Variable** | **Value** | **SE** | **DF** | **t-val** |
| --- | --- | --- | --- | --- |
| (Intercept) | 46.51 | 6.18 | 2012 | 7.53 |
| T_a_ | 0.05 | 0.01 | 2012 | 3.58 |
| Category Shallow Torpor | -3.19 | 0.85 | 53 | -3.74 |
| Category Transition | -16.91 | 0.88 | 53 | -19.31 |
| Category Deep Torpor | -26.60 | 0.77 | 53 | -34.53 |
| Species BLUH | -1.23 | 2.44 | 28 | -0.50 |
| Species RIHU | -2.10 | 2.22 | 28 | -0.95 |
| Capture mass | 0.29 | 0.44 | 28 | 0.67 |
| Year | -0.92 | 0.36 | 28 | -2.59 |
| T_a_ : Category Shallow Torpor | 0.05 | 0.03 | 2012 | 1.67 |
| T_a_ : Category Transition | 0.46 | 0.05 | 2012 | 9.13 |
| T_a_ : Category Deep Torpor | 0.73 | 0.06 | 2012 | 12.69 |
| Category Shallow Torpor : Species BLUH | -0.15 | 0.87 | 53 | -0.17 |
| Category Transition : Species BLUH | 1.74 | 0.98 | 53 | 1.78 |
| Category Deep Torpor : Species BLUH | 1.58 | 0.90 | 53 | 1.75 |
| Category Shallow Torpor : Species RIHU | -0.81 | 0.89 | 53 | -0.91 |
| Category Transition : Species RIHU | 3.25 | 0.86 | 53 | 3.78 |
| Category Deep Torpor : Species RIHU | 2.67 | 0.74 | 53 | 3.59 |

3.	R Core Team. R: A language and environment for statistical computing. R Foundation for Statistical Computing: Vienna, Austria 2018.
