## Supplementary material for "Facultative variation across a shallow to deep torpor spectrum in hummingbirds": Metadata for all datasets

### Introduction

This metadata file describes the data that accompany the hummingbird shallow and deep torpor study conducted June 2017 – June 2018. This study was performed at the Southwestern Research Station in the Chiricahua Mountains in south-eastern Arizona.

### Species table

These species codes are used in some of the data tables below

| *Species code* | *Species scientific name* | *Species common name* |
| --- | --- | --- |
| BCHU | *Archilocus alexandri* | black-chinned hummingbird |
| BLUH | *Lampornis clemenciae* | blue-throated mountain-gem |
| RIHU | *Eugenes fulgens* | Rivoli’s hummingbird |

### Compiled surface temperature maxima

These are the maximum surface temperatures, once per recording (approximately once every 10 minutes) per individual. This file is a “melted” version of the RDS files (see RDS_Data.zip).

**Dataset file**

**Identity:** Thermal_maxes.csv

**Size:** 2104 records, not including header row, 112 kilobytes.

**Format and storage mode:** comma delimited

**Header information:** The first row of the file contains the variable names. See below for detailed descriptions of the column contents

**Alphanumeric attributes:** Mixed

**Special characters/fields:** If no information is available for a given record, or if a value is not appropriate, this is indicated by NA. 0’s indicate true zero.

**Variables**

| *Variable name* | *Variable definition* | *Storage type* | *Variable definitions* |
| --- | --- | --- | --- |
| variable | Represented as Individual ID, underscore, MMDD (month and day) on which the bird was studied | Character | NA |
| value | Maximum surface temperature for that individual for that recording (corresponds to maximum eye surface temperature) | Float | NA |
| Species | Abbreviated species name | Character | See Species table above |
| Indiv_ID | Number of the individual within the species, underscore, MMDDYY (month, day and year) on which the bird was studied | Character | NA |
| Category | Metabolic category that the surface temperature value was assigned to, using the category thresholds assigned. | Character | Levels = Deep Torpor, Normothermic, Shallow Torpor, Transition |

### Assigned metabolic category thresholds per individual

These thresholds were assigned by assessing individual patterns of rates of body temperature change, relative to ambient temperature over the course of the night.

**Dataset file**

**Identity:** Category_thresholds.csv

**Size:** 33 records, not including header row, 2 kilobytes.

**Format and storage mode:** comma delimited

**Header information:** The first row of the file contains the variable names. See below for detailed descriptions of the column contents

**Alphanumeric attributes:** Mixed

**Special characters/fields:** If no information is available for a given record, or if a value is not appropriate, this is indicated by NA. 0’s indicate true zero.

**Variables**

| *Variable name* | *Variable definition* | *Storage type* | *Variable definitions* |
| --- | --- | --- | --- |
| SNo | Row number | Integer | NA |
| Species | Abbreviated species name | Character | See Species table above |
| Individual | Number of the individual within the species, underscore, MMDDYY (month, day and year) on which the bird was studied | Character | NA |
| Normo_min | Lowest temperature which looks to be normothermic for this bird | Float | NA |
| Shallow_max | Highest temp which looks like shallow (same as Normo_min, unless the bird didn’t use shallow torpor, in which case this column has NAs) | Float | NA |
| Shallow_min | Lowest temperature which looks to be shallow torpor for this bird | Float | NA |
| Transition_max | Highest temp which corresponds to transition (same as Shallow_min, whenever both columns are non-NA) | Float | NA |
| Transition_min | Lowest temperature which looks to be in the transition category for this bird | Float | NA |
| Torpor_max | Highest temp which corresponds to deep torpor (same as Transition_min, whenever both columns are non-NA) | Float | NA |

### Interpolated values to ensure evenly spaced, continuous, data

Our raw data were sampled approximately once every 10 minutes. To estimate proportion of the night spent in the different metabolic categories, we needed data points to be evenly spaced. We therefore interpolated the data to get temperature values for every minute. This interpolation function used a linear method to interpolate temperatures based on the two closest temperatures at that time. If the data point was beyond a temperature extreme, the function interpolated using that extreme. We also interpolated the minimum image values to obtain corresponding ambient temperature values for every minute.

**Dataset file**

**Identity:** Interpolated_Thermal.csv

**Size:** 982277 records, not including header row, 66430 kilobytes.

**Format and storage mode:** comma delimited

**Header information:** The first row of the file contains the variable names. See below for detailed descriptions of the column contents

**Alphanumeric attributes:** Mixed

**Special characters/fields:** If no information is available for a given record, or if a value is not appropriate, this is indicated by NA. 0’s indicate true zero.

**Variables**

| *Variable name* | *Variable definition* | *Storage type* | *Variable definitions* |
| --- | --- | --- | --- |
| SNo | Row number | Integer | NA |
| Indiv_pasted | Number of the individual within the species, underscore, MMDDYY (month, day and year) on which the bird was studied | Character | NA |
| Surf_Temp | Interpolated maximum surface temperatures of the bird | Float | NA |
| Amb_Temp | Interpolated ambient temperatures every minute | Float | NA |
| Cap_mass | Capture mass of the individual in grams | Float | NA |
| Category | The four metabolic categories, defined by the thresholds in the Category thresholds file | Character | Levels = Deep Torpor, Normothermic, Shallow Torpor, Transition |
| Time | Military time | Integer | NA |
| Species | Abbreviated species name | Character | See Species table above |

### Percentage of time in each metabolic category

Summarized percentage of time that each species spent in each of the four metabolic categories.

**Dataset file**

**Identity:** Category_percentages.csv

**Size:** 3 records, not including header row, 1 kilobytes.

**Format and storage mode:** comma delimited

**Header information:** The first row of the file contains the variable names. See below for detailed descriptions of the column contents

**Alphanumeric attributes:** Mixed

**Variables**

| *Variable name* | *Variable definition* | *Storage type* | *Variable definitions* |
| --- | --- | --- | --- |
| Species | Abbreviated species name | Character | See Species table above |
| Normothermic | Percentage of time spent normothermic | Float | NA |
| Shallow_torpor | Percentage of time spent in shallow torpor | Float | NA |
| Transition | Percentage of time spent in transition between categories | Float | NA |
| Torpor | Percentage of time spent in deep torpor | Float | NA |

### Bird masses

**Dataset file**

**Identity:** Bird_masses.csv

**Size:** 33 records, not including header row, 2 kilobytes.

**Format and storage mode:** comma delimited

**Header information:** The first row of the file contains the variable names. See below for detailed descriptions of the column contents

**Alphanumeric attributes:** Mixed

**Special characters/fields:** If no information is available for a given record, or if a value is not appropriate, this is indicated by NA. 0’s indicate true zero.

**Variables**

| *Variable name* | *Variable definition* | *Storage type* | *Variable definitions* |
| --- | --- | --- | --- |
| Bird_ID | Original bird ID during data recording | Character | Note: At the time, Rivoli’s hummingbirds were named Magnificent hummingbirds, so RIHU used to be MAHU. |
| Species | Abbreviated species name | Character | See Species table above |
| Day | Day that data collection began | Integer | NA |
| Month | Month of data collection | Integer | NA |
| Month_text | Month stored as text | Character | NA |
| Year | Year of data collection | Float | NA |
| Indiv_ID | Number of the individual within the species, underscore, MMDDYY (month, day and year) on which the bird was studied | Character | NA |
| Capture_mass_g | Bird’s mass at capture, in grams | Float | NA |
| Fed_mass_g | Bird’s fed mass in grams, before it was placed into the chamber for temperature data collection | Float | NA |

### RDS data

This zip file has .rds files, each of which contains all the temperature values for an individual (35 files total).

**Dataset file**

**Identity:** RDS_Data.zip

**Size:** 35 files, 2-4 kilobytes each; 75 kilobytes total when zipped.

**Format and storage mode:** .rds files
